# Kinetics of *de novo* Bone and Bone Marrow Niche Formation with Hybrid Click Cryogels

**DOI:** 10.1101/2025.05.12.653550

**Authors:** Sangmin Lee, Kwasi Adu-Berchie, Azeem Sanjay Sharda, Tania To, Nikolaos Dimitrakakis, Alexander Stafford, Katherine Sheehan, Christopher Johnson, Hamza Ijaz, Phoebe Kwon, Mark Cartwright, Sandy Elmehrath, Mary Catherine Skolfield, Des White, Michael Williams, Michael Super, David T. Scadden, David J. Mooney

**Author notes:** These authors contributed equally: Sangmin Lee, Kwasi Adu-Berchie, Azeem Sanjay Sharda. Corresponding authors., Address: Pierce Hall 319, 29 Oxford Street, Cambridge, MA, 02138.

## Abstract

Successful hematopoietic stem cell transplantation (HSCT) critically depends on efficient T cell recovery, which is limited by compromised bone marrow niches following irradiation. While various factors influence the regeneration of bone and bone marrow niches, the dynamics of this process remain elusive. Here, we explore the kinetics of de novo bone and bone marrow development under varying BMP-2 doses, host immune status, and biological sex, using a cryogel of covalently crosslinked alginate and gelatin releasing BMP-2. Bone formation was monitored by ultrasonography and microcomputed tomography (microCT) analysis, while histological analysis provided insights into the relation between mineralized tissue and bone marrow formation. Bone developed within 2-4 weeks, resulting in cortical bone around the cryogels, and a trabecular bone network with hematopoietic tissue within the cryogels. Higher BMP-2 doses significantly accelerated mineralization kinetics and doubled the resident hematopoietic stem cell population. Notably, immunocompromised status delayed niche development by two weeks and reduced hematopoietic stem cells fourfold. We also found that female mice exhibited enhanced niche formation compared to males under the identical conditions. These findings provide insights into the factors that govern the spatiotemporal regulation of bone and bone marrow niche development and establish this hybrid click cryogel system as a promising platform for improving T cell reconstitution in HSCT patients.

## Introduction

In the treatment of hematological disorders, allogenic hematopoietic stem cell transplantation (HSCT) has become an important therapeutic strategy for severe disease including leukemia and sickle cell anemia ^1–3^. However, this procedure is accompanied by significant immunological challenges and clinical complications. Patients undergoing HSCT are placed in a prolonged state of severe immunodeficiency significantly increasing the risk of infections and complications like Graft-versus-Host disease (GVHD) ^4^. The core of these complex issues lies in the delayed or impaired T cell recovery following bone marrow stress caused by chemotherapy, radiotherapy and the transplantation process ^3^. T cells are integral to the defense mechanism of the immune system, and the recovery of their function is a critical factor in the success of HSCT. Accelerating the reconstitution of immune function is a crucial challenge of this treatment ^5,6^.

Biomaterials may be utilized to artificially generate bone marrow-like microenvironments suitable to house hematopoietic cells and generate T cell progenitors ^7,8^. In particular, the engineering of ectopic bone and bone marrow niches using osteoinductive biomaterials has demonstrated potential to facilitate immune system reconstitution ^9–11^. These strategies encompass various approaches: the use of instructive biomaterials ^12–18^, delivery of osteoinductive cytokines or angiogenetic factors such as bone morphogenetic protein-2 (BMP-2) or vascular endothelial growth factor (VEGF) ^7,19–24^, and genetic modifications ^25^. We have previously demonstrated a methacrylate-alginate cryogel that creates a bone marrow-like niche capable of guiding hematopoietic stem cells (HSCs) toward the T cell lineage and improving T cell reconstitution ^26^. However, the field has limited understanding regarding factors influencing the rate and extent of de novo bone and bone marrow formation, including BMP-2 dose^27^, endogenous factors^28,29^, host immune status ^30^, biological sex ^31–33^ and age^34^.

Here, we explore the impact of BMP-2 dose, biological sex, and immune competence on ectopic bone formation. To better replicate natural bone composition, alginate-gelatin hybrid cryogels were fabricated for BMP-2 delivery, and the kinetics of bone and bone marrow development were investigated through real-time ultrasonography. Data from ultrasonography, microCT analysis, histological analysis, and flow cytometry was compared and correlated to gain a deeper understanding of the dynamics of bone and bone marrow formation. Our findings indicate that the development of bone and bone marrow with hybrid cryogels is largely influenced by the BMP-2 dose. Notably, immunocompromised mice and male mice exhibited delayed and diminished bone and bone marrow-like formation compared to immunocompetent mice and female mice, respectively. Finally, the impact of these biomaterials on post-HSCT immune reconstitution was evaluated.

## Results

### 3.1. Fabrication of hybrid click cryogels

Scaffolds were intended to mimic the structure of bone ^35^, and enable host progenitors to infiltrate and undergo osteogenesis via the release of BMP-2. (**Fig. 1A**) Scaffolds were fabricated by crosslinking tetrazine-modified alginate with norbornene-modified gelatin in a partially frozen state (cryogelation) (**Fig. 1B**). The click reaction between tetrazine and norbornene led to gelation within several minutes **(Supplementary Fig. 1a**) and resulted in cylindrical cryogels containing highly interconnected pores (**Fig. 1C-E**). These cryogels exhibited consistent mechanical properties (**Supplementary Fig. 1b, c**) and maintained high interconnected porosity and shape recovery properties after injection through a 16-gauge needle (**Supplementary Fig. 2**).

**Figure 1.**
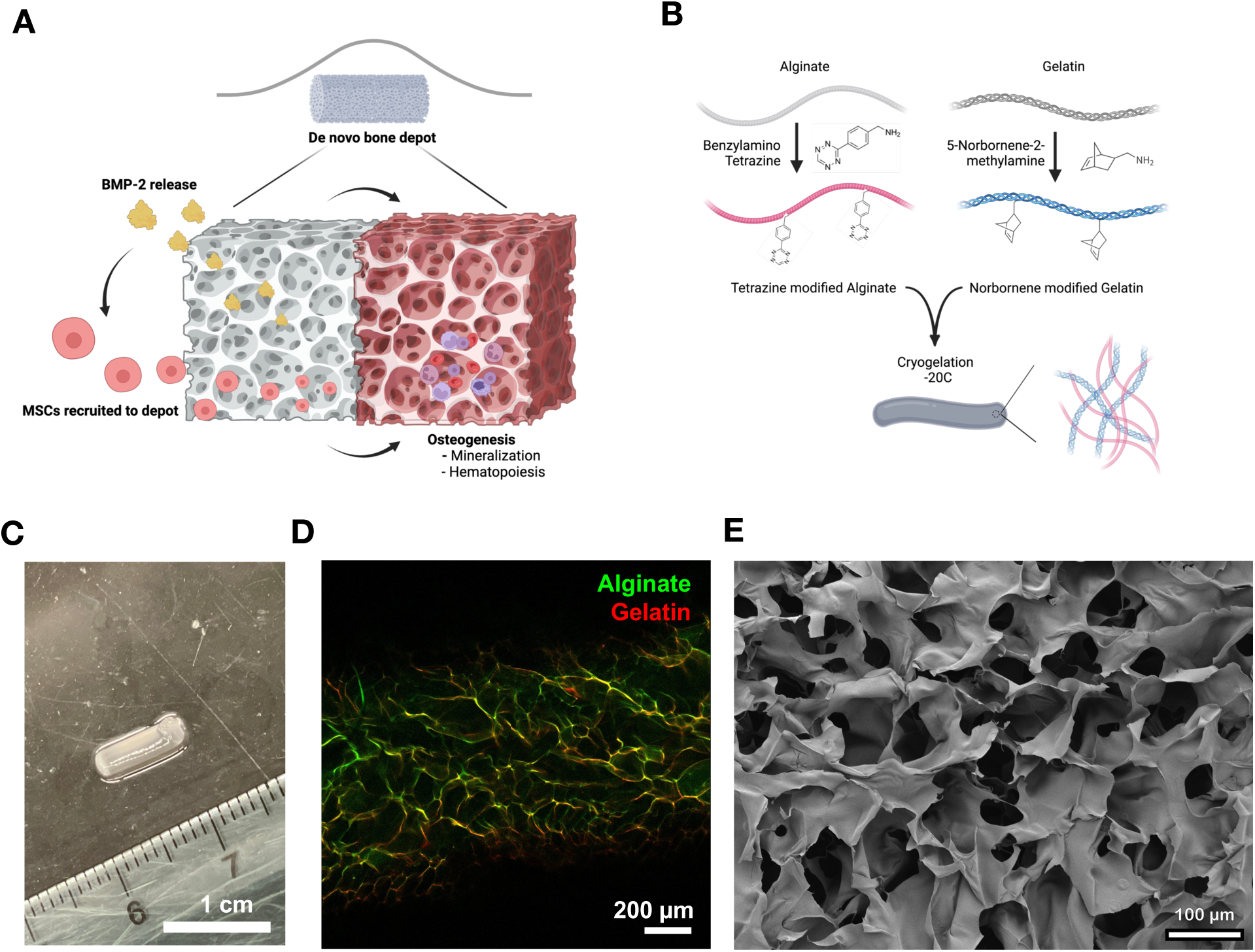
Fabrication and characterization of hybrid click cryogels. (A) Schematic illustrating the concept of hybrid click cryogels for de novo bone formation. (B) Schematic showing the modification of alginate and gelatin with tetrazine and norbornene respectively, and subsequent cryogelation process. (C) Representative optical image of a hybrid click cryogel. (D) High magnification fluorescent image of macroporous hybrid click cryogels. FITC-labeled fluorescent alginate and rhodamine-labeled fluorescent gelatin were used to visualize the polymers in cryogels. (E) High magnification SEM image of hybrid click cryogels.

### 3.2. Kinetics of de novo bone formation from BMP-2 loaded Hybrid Click Cryogels

To investigate bone formation with hybrid click cryogels, BMP-2 was loaded into the cryogels using two approaches: incorporating BMP-2 into the polymer mixture before gelation, and adding BMP-2 post-gelation (**Fig. 2A**). BMP-2, which is positively charged at physiological pH, was expected to bind to both the negatively-charged alginate polymer and negatively-charged amino acid residues in gelatin via electrostatic interactions ^19,36,37^, enabling gradual release. Quantification of BMP-2 release from cryogels (loaded pre-gelation) revealed sustained release over 14 days (**Fig. 2B**). Cryogels containing either 5 ug or 20 ug of BMP-2 were then injected into the subcutaneous tissue of mice, and subsequent mineralization monitored with high-frequency ultrasound sonography (HFUS) (**Fig. 3C**). In ultrasonography images, the cryogels initially appeared grey and hypoechoic relative to the epidermis, while mineralized tissue appeared white and hyperechoic relative to the cryogels. As mineralization progressed and the bone matured, a strong black shadow formed beneath the hyperechoic region. Using these distinct features of bone and cryogels in ultrasonography, we quantified the volumes of gel and mineralization regions over time. Overall, a positive correlation was found between gel volume and mineralization volume. Additionally, the gel volume decreased as mineralization volume increased over time (**Supplementary Fig. 3**). Representative ultrasound images demonstrated strong mineralization of cryogels loaded with 20 ug BMP-2, whereas cryogels with 5 ug BMP-2 exhibited lower mineralization at day 17. By day 31, cryogels with 5 ug BMP-2 demonstrated increased mineralization (**Fig. 2D**). Scoring cryogel mineralization on a 4-point scale (**Supplementary Fig. 4**) revealed that cryogels with 20 ug BMP-2 scored 3 by day 17, increasing to 4 by day 31, while cryogels with 5 ug BMP-2 scored 2 by day 17, reaching 3 by day 31 (**Fig. 2E, F**). No significant differences in mineralization kinetics between pre- and post-gelation BMP-2 incorporation were found.

**Figure 2.**
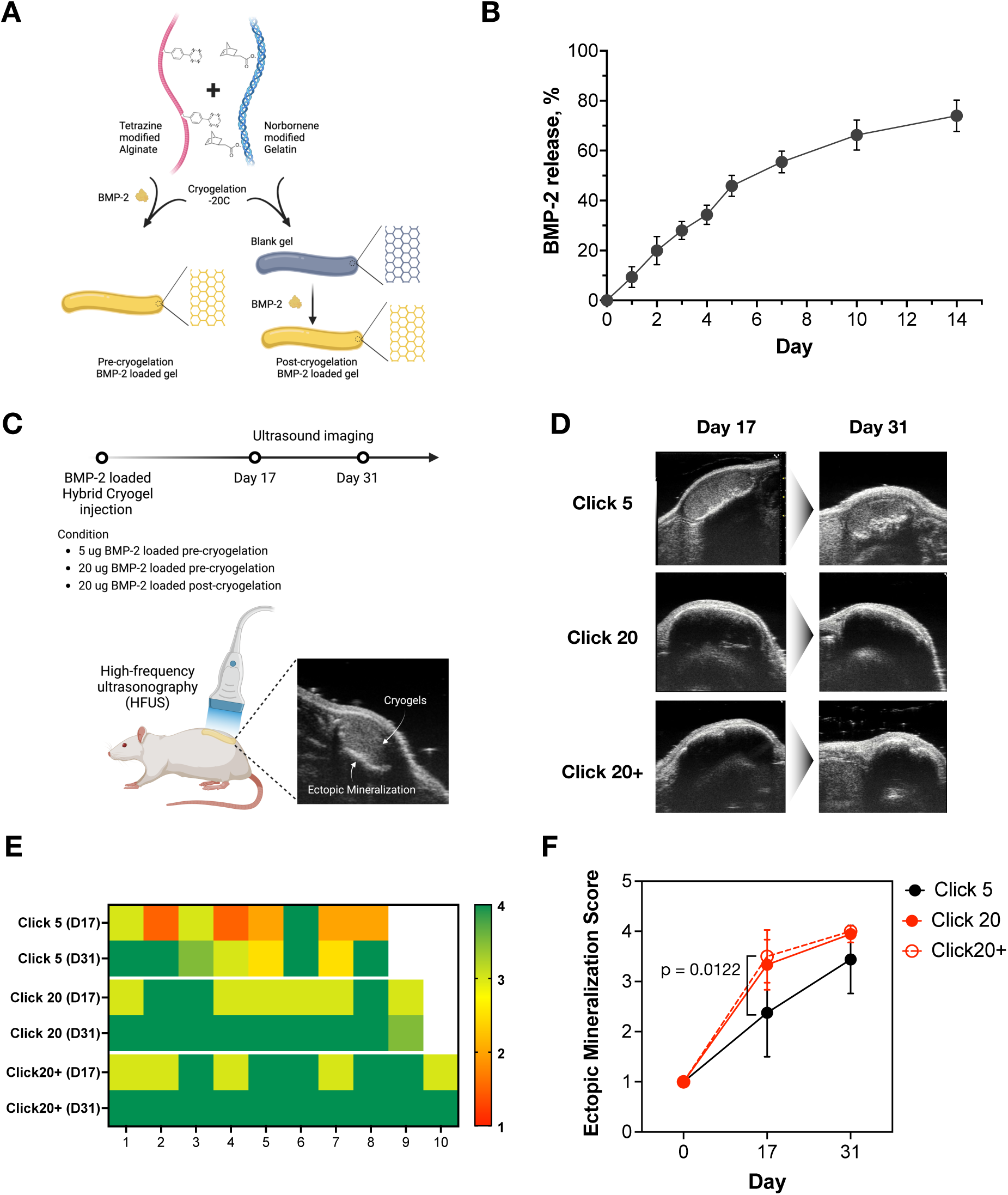
Kinetics of de novo bone formation with hybrid click cryogels using ultrasonography. (A) Schematic comparing BMP-2 loading in cryogels, either during or after cryogelation. (B) BMP-2 release profile from cryogels (BMP-2 loaded during cryogelation). Data represents mean ± standard deviation (n=3). (C) In vivo experimental plan for subcutaneous injection of BMP-2 loaded cryogels, followed by ultrasound imaging using high frequency ultrasonography (HFUS). (D) Representative ultrasonography images demonstrating ectopic mineralization of BMP-2 loaded cryogels at day 17 and day 31. Click 20+ represents condition with BMP-2 loaded post-cryogelation. (E) Ectopic mineralization scores for each individual cryogel for each condition at different time points. (F) Mean ectopic mineralization scores for each cryogel condition over time. Data represents mean ± standard deviation (n=8). An ordinary two-way ANOVA with Tukey’s multiple comparison test was used.

**Figure 3.**
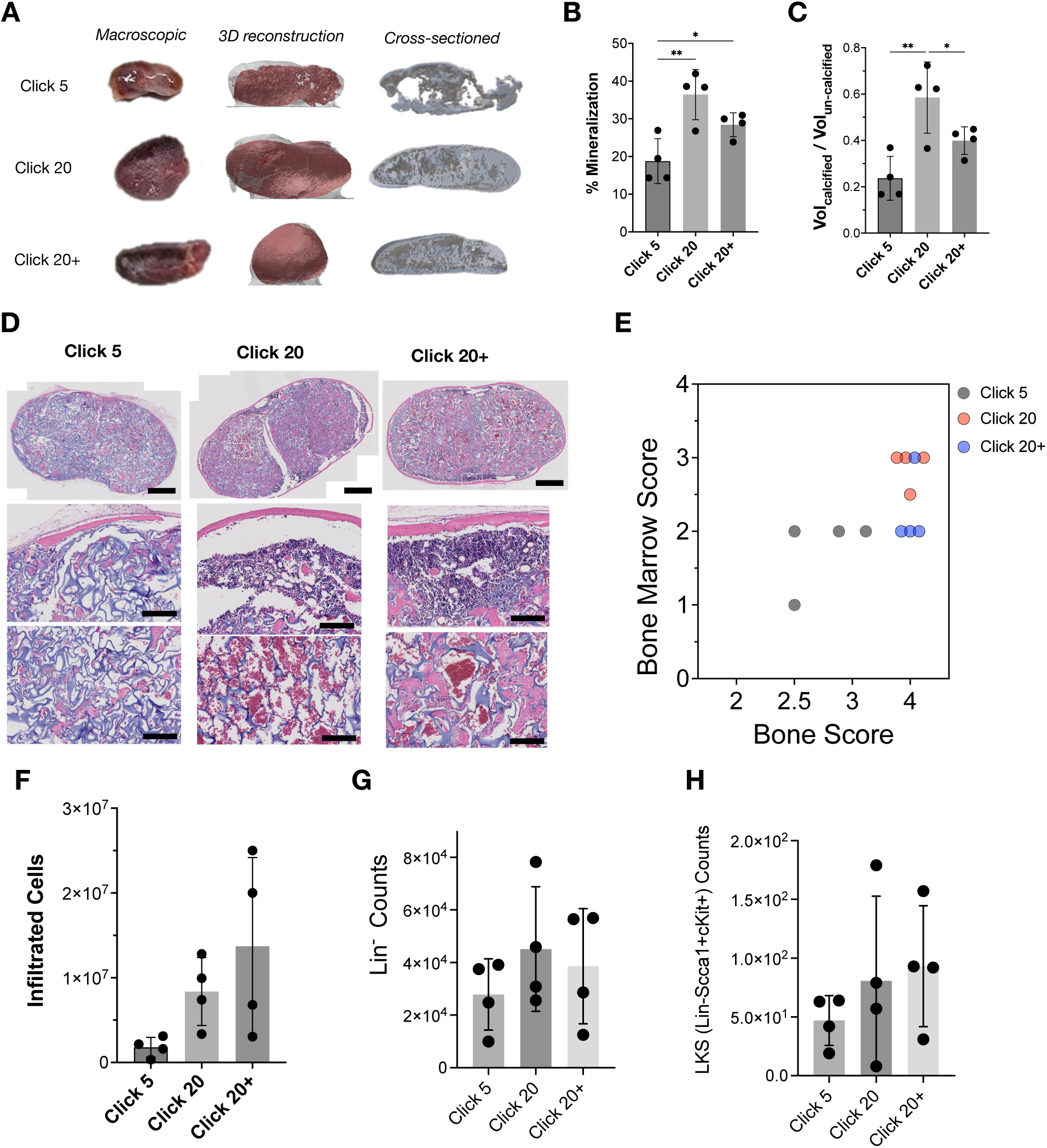
Characterization of de novo bone and bone marrow formation by cryogels. (A) Representative macroscopic optical images, 3D reconstruction and representative tissue cross-sections from microCT analysis of cryogels explanted 35 days post-injection. (B) Percentage of tissue that was mineralized (C), and calcified-to-uncalcified volume ratio of explanted cryogels. Data represents mean ± standard deviation (n=4). An ordinary one-way ANOVA with post hoc Tukey’s multiple comparison test was used; *p <0.05, **p ≤ 0.01. (D) Representative H&E staining of explanted cryogels imaged at low and high magnification. Scale bars represent 500 μm (top row), 100 μm (middle and bottom row). (E) Correlation between bone score and bone marrow score. Flow analysis quantifying the (F) total number of infiltrated cells, (G) lineage-negative cells, and (H) Lin-Sca-1+cKit+(LKS) cells in explanted cryogels. Data represents mean ± standard deviation (n=4).

### 3.3. Impact of BMP-2 dose on bone and bone marrow formation

Cryogels were explanted on day 35 to quantify calcified tissue and bone marrow formation, and cellularity. Macroscopically, the explanted cryogels appeared darker and had a harder texture as compared to cryogels at the time of implantation, and cryogels loaded with 20 ug BMP-2 appeared larger than those with 5 ug BMP-2 (**Supplementary Fig. 5**). Qualitatively, microCT analysis revealed newly developed bone covered the entire gel, forming a trabecular-like network, in cryogels with 20 ug BMP-2. In contrast, cryogels loaded with 5 ug BMP-2 exhibited a discontinuous shell of mineralized tissue with large pores (**Fig. 3A**). Quantification confirmed that cryogels with 20 ug BMP-2 had greater mineralization, a higher calcified-to-uncalcified volume ratio and thicker trabeculae compared to those with 5 ug BMP-2. Additionally, pre-incorporated of BMP-2 (20 ug dose) led to higher calcified-to-uncalcified volume ratio and thicker trabeculae than those with post-incorporated BMP-2 (**Fig. 3B, C, Supplementary Fig. 6**). Comparison of total cryogel volumes estimated in situ via ultrasound imaging and after dissection with microCT analysis revealed a linear correlation between the two measurements. Additionally, comparison of bone score based on ultrasound imaging and calcified volume after dissection with microCT analysis revealed a linear correlation between the two measurements (**Supplementary Fig. 7**).

The distribution of bone and bone marrow in the cryogels was next analyzed through histological staining with H&E and Masson’s trichrome. In the 20 ug BMP-2 gels, bone formed predominantly at the cryogel edges, with spicules extending to the interior, while bone marrow hematopoietic niches with high cell densities were present near the gel periphery. Red blood cells (RBCs) were apparent throughout the gel volume. In contrast, the 5 ug BMP-2 gels exhibited a disconnected bony shell with small spicules within the cryogels, alongside marginal hematopoietic niches and minimal RBC infiltration (**Fig. 3D, Supplementary Fig. 8**). The quantities of mineralized tissue and bone marrow were then analyzed from H&E-stained images using a 4-point scale (**Supplementary Fig. 9a, b**). The mineralized tissue scores positively correlated with the presence of bone marrow across all conditions, with higher scores observed for both types of 20 ug BMP-2 gels compared to 5 ug BMP-2. Notably, pre-incorporated BMP-2 gels exhibited greater mineralized tissue and bone marrow formation than post-incorporated BMP-2, consistent with microCT findings (**Fig. 3E, Supplementary Fig. 9c, d**).

The cell populations in the formed bony tissues were next analyzed through flow cytometry. The 20 ug BMP-2 gels contained 10 times more infiltrated cells than the 5 ug BMP-2 gels (**Fig. 3F, Supplementary Fig. 10a**). Among these, 70% were lineage-negative cells, indicating that the cell populations near the peripheral bone regions likely comprise a hematopoietic niche. Further analysis revealed 0.2% of lineage negative cells were hematopoietic stem and progenitor cells (Lin-Sca1+cKit+). Although the 20 ug BMP-2 gels showed a slightly higher hematopoietic cell population than the 5 ug BMP-2 gels, the increase was not statistically significant (**Fig. 3G, H and Supplementary Fig. 10b, c**). Mature T cells, myeloid cells, and B cells were also present in the gels (**Supplementary Fig. 10d**).

### 3.4. Impact of biological sex and immunocompetency on bone formation

Next, we investigated how biological sex and immunocompetency of mice affected the kinetics of de novo bone formation from BMP-2 loaded cryogels. Cryogels with 5 and 20 ug BMP-2 were injected subcutaneously in male and female mice with lethally irradiation or not. We designated the lethally irradiated mice as having an immunocompromised condition and the intact mice as having an immunocompetent condition. Immunocompromised mice received HSC transplantation in addition to cryogel injection. De novo bone formation was tracked and scored via ultrasonography on days 17 and 33 (**Fig. 4A**). Immunocompetent mice exhibited faster and more extensive mineralization than immunocompromised mice especially in the 5 ug BMP-2 group, while 20 ug BMP-2 group show similar mineralization kinetics. This difference was gradually diminished over time in 5 ug BMP-2 group (**Fig. 4B, C**). In both immunocompromised and immunocompetent groups, male mice showed slower and reduced mineralization compared to female mice at the 5 ug BMP-2 dose. However, as the BMP-2 dose increased, mineralization differences between male and female mice diminished, with no significant sex-based differences observed, regardless of immunocompetency at the 20 ug dose. (**Fig. 4D, E**)

**Figure 4.**
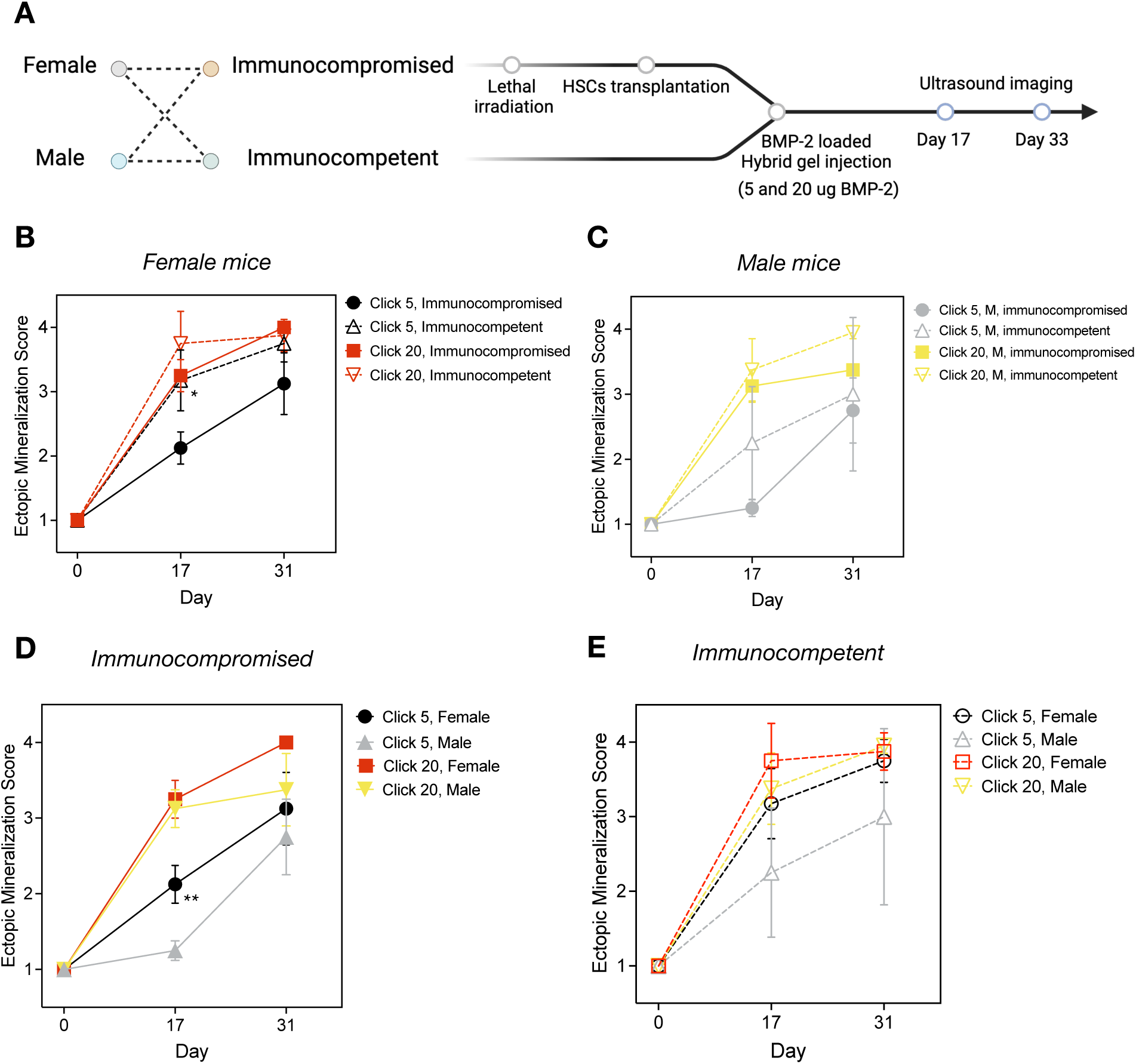
Effect of biological sex and immunocompetency on de novo bone formation kinetics. (A) In vivo experimental plan for subcutaneous injection of BMP-2 loaded cryogels in either female or male, immunocompromised or immunocompetent mice, followed by ultrasound imaging using HFUS. Comparison of de novo bone formation kinetics between immunocompromised and immunocompetent conditions in (B) female mice and (C) male mice. Comparison of de novo bone formation kinetics between female and male mice in (D) immunocompromised and (E) immunocompetent condition. Data represents mean ± standard deviation (n=4). An ordinary one-way ANOVA with post hoc Tuckey’s multiple comparison test was used; *P <0.05.

The cryogels were extracted on day 35 to further assess the amount and quality of de novo bone formation. Consistent with ultrasonography findings, microCT revealed that female mice exhibited relatively greater mineralization than male mice in the 5 ug BMP-2 group, regardless of immunocompetency. In the 20 ug BMP-2 group, no significant differences were observed between male and female mice within the same immunocompetency group, nor between immunocompromised and immunocompetent mice within the same sex group (**Fig. 5A, B, Supplementary Fig. 11**). Further analysis scored the quality of bone and bone marrow from histology results. Immunocompetent mice exhibited pronounced cortical bone with spicules (bone score: 4) and had medium to large size region of hematopoiesis, either at the periphery or inside the scaffold (bone marrow score: 3 or 4) in both sexes, while immunocompromised mice, particularly males, showed robust bone formation but had smaller bone marrow niches in the peripheral region (bone marrow score: 2) or no bone marrow (bone marrow score:1) (**Fig. 5C, D**). In the immunocompetent group, both females and males developed substantial bone and bone marrow niches in the 20 ug BMP-2 group. In the immunocompromised group, bone formation was prominent in both sexes, but bone marrow was more prominent in females than males, and within the 20 ug BMP-2 group. (**Fig. 5E, F**) The quantities of lineage negative cells in explanted cryogels mirrored the results seen in bone and bone marrow scores. The immunocompetent group showed a greater number of lineage negative cells compared to the immunocompromised group. Interestingly, while the 20 ug BMP-2 dose resulted in a relatively higher number of cells compared to 5 ug BMP-2 in the immunocompromised group, similar numbers of lineage negative cells were observed for both 5 ug and 20 ug BMP-2 treatments in the immunocompetent group, regardless of sex (**Fig. 5G, H**). The number of LKS (Lin-cKit+Sca1+) cells measured on day 35 after irradiation was BMP-2 dose dependent in the immunocompromised group, but not in the immunocompetent group **(Supplementary Fig. 12, 13**).

**Figure 5.**
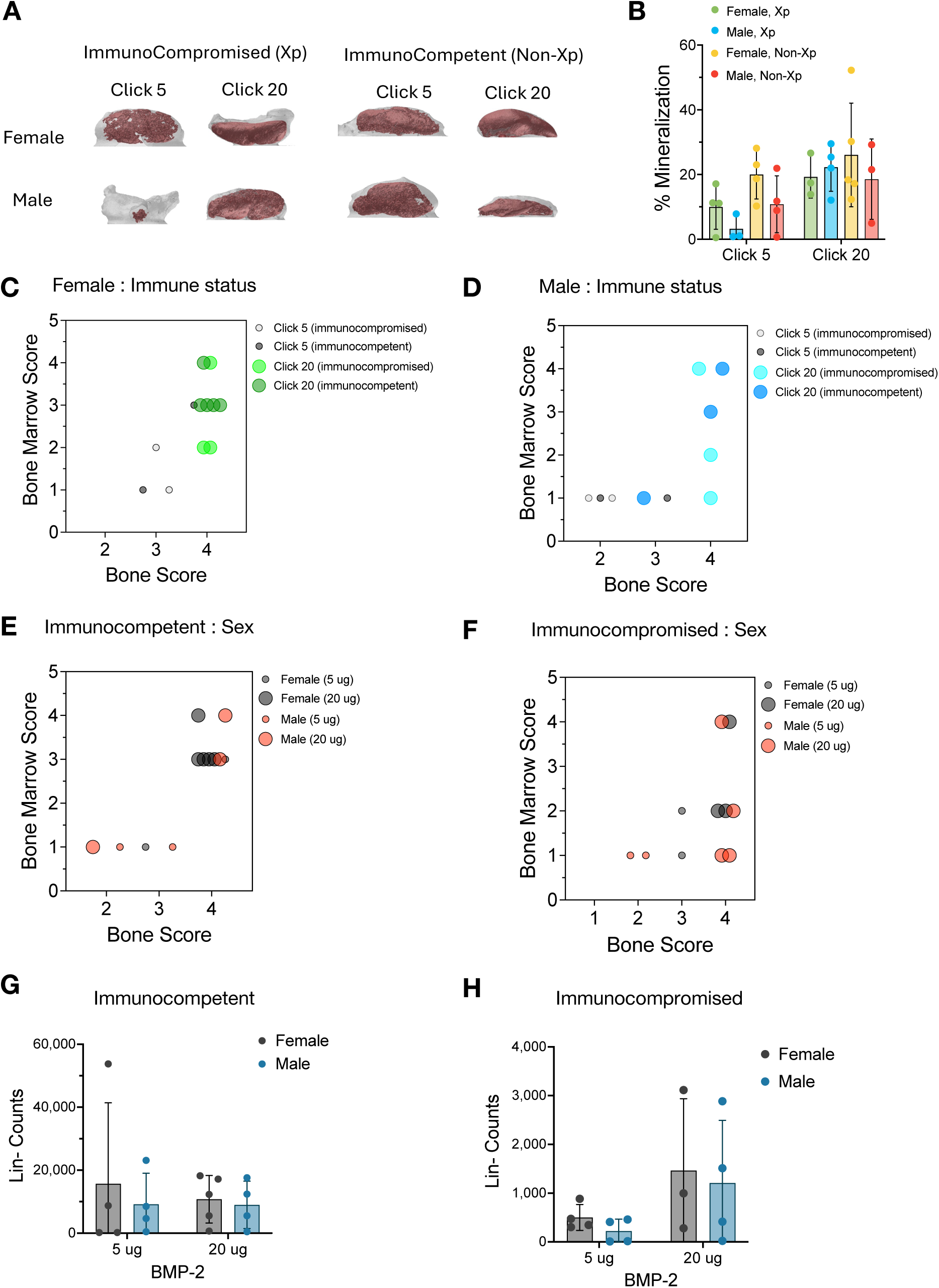
Characterization of de novo bone formation amount and quality as a function of sex and immunocompetency. (A) Representative microCT images of explanted cryogels (day 35). (B) Percentage of mineralization in explanted cryogels. (C) Histological analysis-based bone and bone marrow scores in (C) female mice and (D) male mice under different immune statuses: immunocompromised or immunocompetent. (E) Histological analysis-based bone and bone marrow scores in (E) immunocompetent and (F) immunocompromised conditions under different sex. Data represents mean ± standard deviation (n=4). Flow analysis quantifying the population of lineage-negative cells in (G) immunocompetent and (H) immunocompromised conditions. Data represents mean ± standard deviation for n=4 cryogels.

### 3.5. T cell reconstitution from ectopic bone

We next examined the impact of cryogel-mediated bone formation on the kinetics and magnitude of T cell reconstitution following lethal irradiation and hematopoietic stem cell (HSC) transplantation. Lethally irradiated, HSC-transplanted mice were injected with either blank click hybrid cryogels or click hybrid cryogels loaded with 20 ug BMP-2 (**Fig. 6A**). As a control, we examined the peripheral blood analysis of mice without cryogels, with or without lethal-irradiation, and it showed the significantly decreased T cell numbers in immunocompromised conditions compared to immunocompetent conditions **(Supplementary Fig. 14).** Sequential blood analysis at various time points after cryogel injection revealed that mice receiving BMP-2 loaded cryogels showed a trend for faster reconstitution kinetics for white blood cells, lymphocytes, and donor T cells compared to those receiving blank cryogels in female mice (**Fig. 6B, C, Supplementary Fig. 15**). Also, females showed significantly larger donor T cell populations after 6 weeks of transplantation compared to males, while there was no significant difference in B cell populations over time (**Supplementary Fig. 16a**). After 84 days, the cryogels were explanted to examine the cellular populations within the BMP-2 loaded cryogels and compare to those in blank cryogels. The number of infiltrated cells in BMP-2 loaded cryogels was over 100 times higher than in blank cryogels. Significantly higher numbers of donor T cells, lineage-negative cells, and LKS cell populations were found residing in the BMP-2-loaded cryogels compared to the blank cryogels. This suggest that the HSCs circulating in the blood proliferated and differentiated within the BMP-2-loaded cryogels (**Fig. 6D-G**). Moreover, females showed significantly larger populations of immune cells including T cells, B cells, and neutrophils, as well as hematopoietic cells including lineage-negative cells and LKS cells, in the BMP-2-loaded cryogels compared to males (**Supplementary Fig. 16b**). This trend of females outperforming males in terms of immune cell and hematopoietic cell population development in cryogels was only noted at this later time point, as we observed no significant difference between females and males at day 35 (**Supplementary Fig. 12b, c**). Histology results showed distinct bone marrow-like hematopoietic tissue residing near the ectopic bone in BMP-2 loaded cryogels, whereas cell infiltration in blank cryogels was minimal (**Fig. 6H, I**). In the bone marrow-like niches, the majority of cells expressed CD45, indicating hematopoietic lineage, while fewer cells infiltrating blank cryogels expressed CD45. Notably, some cells within the bone marrow-like niches also expressed Notch intracellular domains (NICD) and DLL4 (**Fig. 6J, Supplementary Fig. 17**), signaling pathways that drive the differentiation of HSCs into T cell-lineage populations.

**Figure 6.**
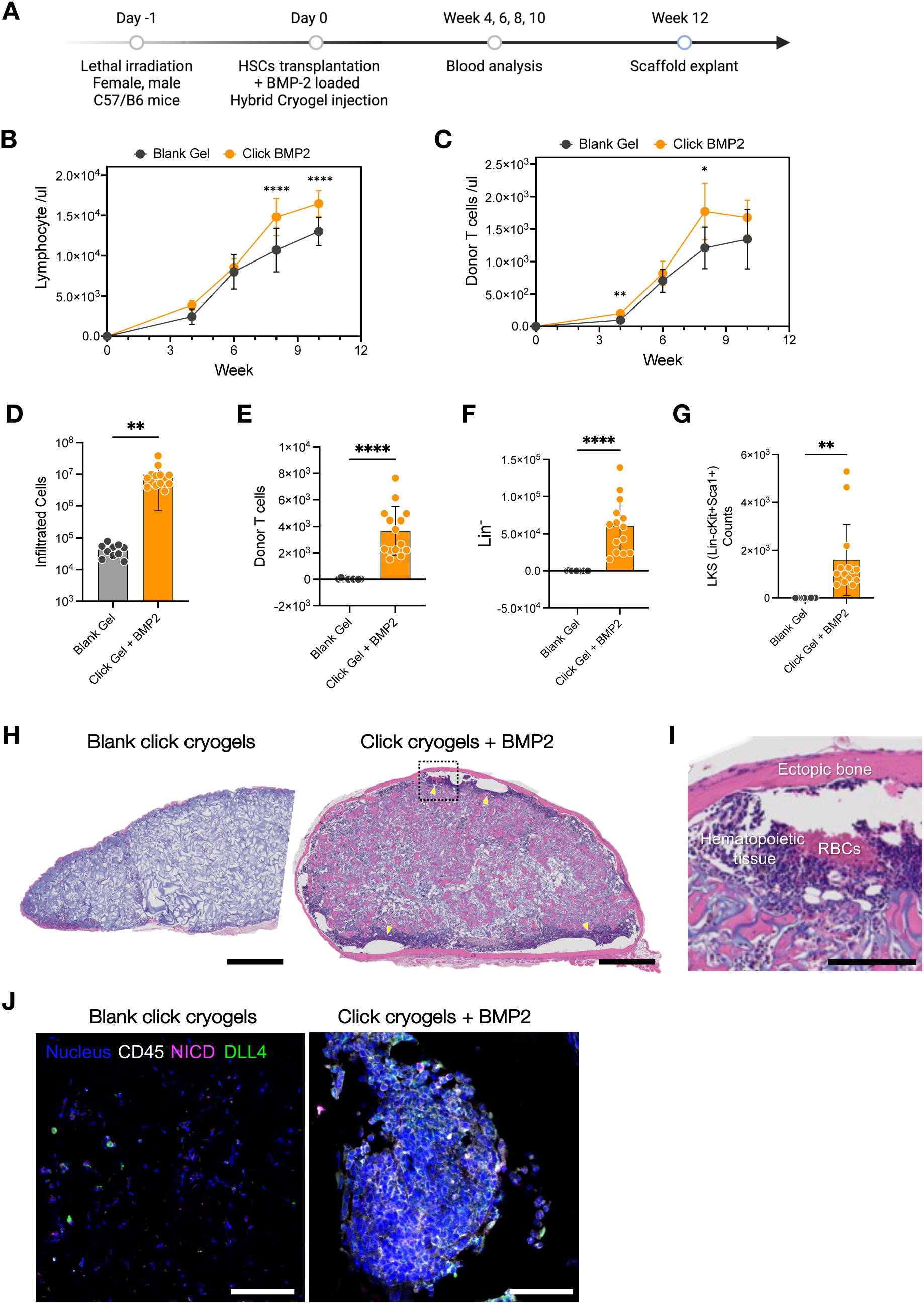
T cell reconstitution from ectopic bone. (A) In vivo experimental plan for T cell reconstitution through lethal irradiation and HSCs transplantation, followed by subcutaneous injection of BMP-2 loaded cryogels. Quantification of circulating immune cell population: (B) lymphocytes and (C) donor T cells over time. Data represents mean ± standard deviation (n=10). An ordinary two-way ANOVA with Šídák’s multiple comparison test was used; *p ≤ 0.05, **p ≤ 0.01, ****p ≤ 0.0001. (D-G) Cell counts in explanted cryogels: (D) infiltrated live cells, (E) donor T cells, (F) lineage-negative cells, and (G) LKS cell population. Data represents mean ± standard deviation (n=10). An unpaired t-tests using Welch’s correction was used; **p ≤ 0.01, ****p ≤ 0.0001. (H-I) Representative H&E images of blank cryogels and BMP-2 loaded click hybrid cryogels at (H) low and (I) high magnification. Yellow arrows depict the hematopoietic tissue region. Scale bars represent 500 μm (H) and 100 μm (I). (J) Representative immunohistochemistry images of explanted blank cryogels and BMP-2 loaded click hybrid cryogels, showing staining for different cell populations: immune cells in white, Notch intracellular domains (NICD; Notch-activated cells) in pink, DLL4-expressing cells in green. (Nucleus: blue; CD45+: white; NICD: pink; DLL4: green) Scale bars represent 50 μm.

## Discussion

This study explored the kinetics of de novo bone and bone marrow development under varying BMP-2 doses, host immune status, and biological sex using BMP-2 loaded hybrid click cryogels. Multimodal analysis, including ultrasonography, microCT, histology, and flow cytometry, revealed a linear correlation between bone and bone marrow-like niche development in biomaterials-based platform. These findings revealed that higher BMP-2 doses, immunocompetency, and female mice resulted in significantly faster mineralization and hematopoiesis with a larger population of hematopoietic stem/progenitor cells compared to lower BMP-2 doses, immunocompromised conditions, and male mice. Finally, BMP-2 loaded hybrid click cryogels expanded hematopoietic stem/progenitor cells and accelerated the expansion of donor T cells compared to blank cryogels.

The biomaterial strategy developed here addresses limitations of the traditional biomaterial-based platforms used for ectopic mineralization - disk-shaped implantable scaffolds or injectable in situ forming gels. While implantable scaffolds provide reliable mechanical support, they necessitate invasive surgeries with associated inflammation ^10^. Injectable in situ forming gels offer a minimally invasive alternative and can be effective when an injection pocket is available. However, these gels often lack mechanical stability due to a limited window for gelation time and concentration constraints necessary for injectability. The injectable, rod-shaped hybrid click cryogels described in this study address these limitations, as these pre-formed scaffolds utilize click chemistry to provide precise control over mechanical properties while maintaining injectability through standard syringes, enabling minimally invasive delivery.

The hybrid click cryogels we developed serve as a platform for ectopic bone and bone marrow-like niche development, with their macroporous structure providing high surface area for effective BMP-2 loading and controlled release. We loaded BMP-2 into hybrid click cryogels through two different approaches: during cryo-gelation or after cryo-gelation. Our observations suggest that the cryo-gelation process facilitates more effective binding between BMP-2 and alginate. This enhanced binding may be attributed to the space confinement of BMP-2 in the polymer-rich phase during cryo-gelation, which appears to accelerate binding compared to the diffusion-derived binding that occurs when loading BMP-2 after cryo-gelation. Moreover, we observed slightly better mineralization and hematopoietic tissue formation when BMP-2 was loaded during cryo-gelation, indicating that this method may be preferable for tissue engineering outcomes. Our results demonstrate that BMP-2 loaded hybrid click cryogels successfully recapitulate key features of native bone tissue, including both cortical and trabecular bone structure, along with hematopoietic tissue resembling the bone marrow niche, in a dosage-dependent manner. Higher BMP-2 doses not only accelerated the mineralization process but also resulted in more extensive hematopoietic tissue formation, with a twofold increase in the hematopoietic stem cell population compared to lower doses. These observations align with previous studies that have established BMP-2 as an osteoinductive cytokine that promotes mineralization ^38–40^. Here, notably, BMP-2 maintains its biological activity despite exposure to partial freezing during the cryo-gelation process.

The multimodal analysis of de novo bone and bone marrow-like niche formation in hybrid click cryogels revealed a linear correlation between mineralized tissue and bone marrow development. While microCT analysis has been traditionally used to analyze mineralization quantitively, it lacks the ability to quantify hematopoietic tissue development ^41–43^. To address this limitation, we introduced a novel histology data-based scoring metric that can assess the extent of both bone formation and bone marrow development within the same sample. Validating our approach, we found that the bone scores correlated strongly with microCT quantification data (**Supplementary Fig. 7c**), while bone marrow scores showed consistent trends with flow cytometry analyses of hematopoietic cell population. Further, the kinetics of mineralization significantly affected bone and bone marrow development, as demonstrated through in-situ ultrasonographic monitoring and histology analysis of hybrid click cryogels. Notably, slower mineralization rates resulted in reduced hematopoietic tissue formation compared to faster mineralization processes, despite both conditions achieving similar extent of mineralization. This observation suggests that the rate of mineralization, not just final mineral content, plays a crucial role in determining the development of hematopoietic tissue.

Our investigation into de novo bone and bone marrow development reveals critical differences in these processes between immunocompromised and immunocompetent animals. In immunocompromised mice, induced by irradiation and hematopoietic stem cell transplantation, we observed delayed mineralization and reduced hematopoietic tissue development compared to immunocompetent mice, consistent with previous studies demonstrating compromised bone quality in the absence of a functional host immune system, especially mature T and B cells ^10,44^ (**Supplementary Fig. 14)**. While this impairment was evident across BMP-2 doses, higher doses partially mitigated these effects, suggesting a dose-dependent compensatory mechanism on immunodeficiency. To understand the underlying mechanisms, we examined cellular populations and found significantly decreased T and B cell numbers in immunocompromised conditions (**Supplementary Fig. 13, 14**). This depletion of immune cells, along with the reduction in stromal cells and vascularization due to irradiation, likely directly impacts the BMP-2-induced ectopic mineralization process. Typically, the process of de novo bone formation begins with stromal cell recruitment followed by osteogenesis ^10,45,46^. However, our results, consistent with prior research ^47–50^, suggests that irradiated stromal cells were less effective in activating and differentiating into osteoblasts during BMP-2-induced ectopic mineralization compared to intact stromal cells. The effects of the immunocompromised condition extended beyond bone formation to the hematopoietic niche. We observed significant decreases in lineage-negative cells and hematopoietic stem cells (Lin-Sca1+c-kit+) within mineralized cryogels, particularly at lower BMP-2 doses (**Supplementary Fig. 13**).

Biological sex was found to play an important role in de novo bone and bone marrow development under immunocompromised conditions. Our findings indicate that male mice exhibited slower mineralization kinetics and developed smaller hematopoietic tissues compared to females with the same BMP-2 dosage. These results challenge previous studies that have shown females are generally more compromised in bone development due to their hormones and higher osteoclast resorptive activity compared to males ^51–54^; however, it is important to note that most of the previous studies were conducted under homeostatic conditions. In our study, we observed that this sexual effect becomes less pronounced in immunocompetent animals, suggesting that the inflammatory state plays a role in modulating sex-based differences in bone development. We hypothesize that sex differences in immune response between females and males contribute to difference in bone formation, at least in part. This hypothesis is supported by our observation of higher neutrophil levels in females under immunocompromised conditions (**Supplementary Fig. 12**), along with previous studies showing that females generally have stronger immune responses than males ^55–57^, which may explain the enhanced bone development we observed in females. Furthermore, we find superior T cell reconstitution in females, consistent with the findings on bone development.

In conclusion, these findings demonstrate the efficacy of injectable, rod-shaped hybrid click cryogels as a versatile platform for ectopic bone and bone marrow-like niche formation. This approach overcomes certain limitations associated with previous biomaterial-based platforms. The BMP-2 loaded cryogels promoted formation of tissues with features of native bone in a dose-dependent manner. Our findings highlight the critical role of mineralization kinetics, host immune system, and biological sex in de novo bone and bone marrow development within the context of hybrid click cryogels. These results not only advance our understanding of ectopic bone formation but also suggest potential therapeutic applications, particularly in enhancing T cell reconstitution following HSCT.

## Methods

### Chemical modification of Alginate tetrazine and Gelatin norbornene

Pronova ultrapure MVG sodium alginate (CAS: 9005-38-3, NovaMatrix, Cat# 4200101) was modified with 3-(p-benzylamino)-1,2,4,5-tetrazine (Tz) using carbodiimide chemistry. Briefly, alginate was first dissolved in 0.1 M 2-(N-morpholino) ethanesulfonic acid (MES), 0.3 M NaCl, pH 6.5 at 10 mg/ml. Next, 3.7 g of 1-ethyl-3-(3-dimethylaminopropyl)-carbodiimide hydrochloride (EDC) (ThermoFisher #22980) and 2.2 g of N-hydroxysuccinimide (NHS) (ThermoFisher #24500) were added per gram of alginate. 0.2 g of tetrazine was then added to the dissolved alginate under constant stirring and allowed to react overnight at room temperature. The tetrazine-modified alginate was subsequently purified first by centrifugation at 11,000xg for 15 minutes at room temperature, followed by tangential flow filtration (KrosFlow KR2i; Spectrum Labs) against a 150 to 0 mM decreasing NaCl gradient using a 3kDa MWCO membrane (Repligen, Cat# S02-E003-05-N), and then by treatment with 1 g activated charcoal. This was followed by filtration through a 0.22-µm filter and subsequent lyophilization.

Fish Gelatin from cold water fish skin (Sigma Cat #G7041) was modified with 5-Norbornene-2-methylamine (Nb) using carbodiimide chemistry as follows. Briefly, 1g gelatin was dissolved at a final concentration of 5mg/ml in 0.1M MES, 0.3M NaCl buffer, pH 6.5 on a hotplate heated to 37°C. Next, 0.38g of EDC and 0.115g of NHS were added per gram of gelatin. 123ul of Nb was then per gram of gelatin under constant stirring and reacted at 37°C overnight. The solution was then dialyzed against decreasing NaCl gradient (150mM to 0mM) for 3 d and lyophilized.

### Preparation of Hybrid Click Cryogels

To fabricate 2 % wt/vol alginate–gelatin hybrid cryogels comprising 45% alginate and 55% gelatin, lyophilized Gelatin Norbornene (Gel-Nb) was first dissolved to 10 mg/ml in milliQ water and cooled to 4°C. Alginate Tetrazine (Alg-Tz) was freshly dissolved to 2% wt/vol in milliQ water and also cooled to 4°C. Dissolved Gel-Nb and Alg-Tz were mixed, yielding final concentration of 1.1 mg/ml and 0.9 mg/ml for gelatin and alginate, respectively. The cryogel mixture was immediately pipetted into pre-chilled 2 mm diameter Tygon tubing (VWR #89404-3018) at 100 ul cryogel mixture per 2 cm tubing, and placed in a −20°C freezer overnight for cryo-polymerization. After cryogelation, the cryogels were thawed at room temperature and ejected by gently flushing the tygon tubing with media (DMEM, high glucose, Glutamax; Gibco, #10566-016).

Recombinant Human/Mouse/Rat BMP-2 (R&D systems #355-BM/CF) was incorporated into cryogels by either adding them to the gel mixture and underwent cryogelation as described above or adding them after cryogelation by submerging the hybrid click cryogels into a concentrated BMP-2 solution for 3 hours at 4°C.

The BMP-2 release from the hybrid click cryogels was measured using enzyme-linked immunosorbent assay (ELISA; R&D systems, DY355). The BMP-2 loaded hybrid click cryogels were incubated in media (DMEM, high glucose, GlutaMAX + 10% FBS + 1% pen/strep) at 37 °C. The supernatants were collected at each timepoint and replaced with fresh media. Collected supernatants were stored at −80°C until the ELISA was performed.

### Characterization of Hybrid Click Cryogels

#### Scanning Electron Microscopy (SEM)

Cryogels were first fragmented cross-sectionally to expose the inner surface area. Fragmented gels were serially dehydrated by submersion into increasing concentrations of ethanol from 50% to 100%. The samples were further dried using liquid CO2 in the Tousimis 931 GL critical point dryer. Dried cryogels were mounted onto carbon tape on sample chucks. 10 nm of platinum-palladium was deposited on the sample surface using the EMS 150T ES Sputter/Carbon Coater. Samples were imaged with the FESEM (Zeiss Gemini 260 field emission scanning electron microscope) at 3 kV with 7 mm working distance and were taken using the secondary emission (SE2) detector.

### Mechanical properties

Shear rheological analysis was performed on the gels as follows. Lyophilized alginate tetrazine and gelatin norbornene polymers were first dissolved in milliQ water, cooled to 4C and combined in a pre-cooled Eppendorf tube, resulting in 1.1 mg/ml and 0.9 mg/ml final concentrations of gelatin and alginate respectively. 700ul of the gel solution was immediately cast onto a pre-cooled Peltier plate of a CMT rheometer (AR-G2, TA Instrument). A 40mm cone geometry was then used to perform oscillatory time sweep test at 25°C for 2 hours at a 1% strain and 1Hz frequency.

Compressive elastic modulus analysis was performed on the pre-formed cryogel. Uniaxial compression tests were conducted using a Bose machine to determine the elastic modulus of the hybrid click cryogels. The cryogels were placed on top of a holder and wicked to remove excess water to prevent slippage during the testing. The cryogels were compressed at a rate of 0.1 mm/min, and the elastic modulus was calculated as the slope of the stress-strain curve between 0 and 15% strain.

### Estimation of cryogel interconnected porosity and shape recovery

Cryogel interconnected porosity and shape recovery were estimated as previously described ^58^. Briefly, interconnected porosity was estimated by first weighing intact cryogels (M_hydrated_) and then wicking cryogels with a kimwipe for 15 s to remove the excess buffer from cryogel pores (M_wicked_). Interconnected porosity was quantified as follows:

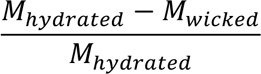

Equation (1): Estimation of interconnected porosity

Cryogel shape recovery was estimated by first weighing intact cryogels (M_hydrated_) and wicking as described above. Wicked cryogels were then rehydrated in DPBS and weighed (M_rehydrated_). Shape recovery was quantified as follows:

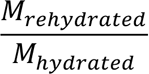

Equation (2): Estimation of cryogel shape recovery

### Animal studies

All animal work was approved by the Harvard Institutional Animal Care and Use Committee (IACUC) and followed the National Institutes of Health guidelines and relevant ethical regulations. C57BL/6 (B6, H-2b), C57BL/6 (CD45.1+) and CByJ.B6-Tg(UBC-GFP)30Scha/J (GFP) (Jackson Laboratories) were females or males and between 6 and 8 weeks old at the start of the experiments. All mice in each experiment were age matched and no randomization was performed.

#### In vivo ectopic bone and bone marrow development study

To investigate ectopic bone and bone marrow development in vivo, BMP-2-loaded hybrid click cryogels were subcutaneously injected into the flanks of female and male C57BL6/J (B6) and C57BL/6 (CD45.1+) mice (Jackson Laboratories) aged between 6 and 8 weeks. Mice were anesthetized when they received subcutaneous injections of two hybrid click cryogels, one in each flank (right and left), which were suspended in saline solution and administered through a 16-gauge needle.

#### In vivo ectopic bone and bone marrow at immunocompromised study

To recapitulate the allo-HSCT procedures and create immunocompromised conditions, lineage-depleted bone marrow cells were isolated from C57BL/6 (CD45.1+) mice. All total-body irradiation procedures were performed using a cesium-137 gamma-radiation source with a single dose of 1,000 cGy on C57BL/6 (B6) mice. Subsequently, 5 x 10^5^ lineage-depleted GFP+ bone marrow cells were transplanted via retro-orbital injection, and BMP-2-loaded hybrid click cryogels were subcutaneously injected. *T cell reconstitution study*: all cohorts of mice were serially bled: bi-monthly, starting 4 weeks post-transplant. White blood cells, lymphocytes, donor T cells, B cell were quantified by CBC (complete blood count) analysis using an Abaxis VetScan HM5.

### Ultrasound imaging and analysis

For ultrasound imaging, mice were anaesthetized and placed on a heating pad with their limbs fixed to prevent movement. Subcutaneously-injected hybrid click cryogels were analyzed using a high-frequency ultrasound device (Vevolab 3100, FUJIFILM VisualSonics) and imaged using a Vevo 3100 scanner with a 50 MHz transducer (MX700) and a 3D motor (VisualSonics; axial resolution, 30 μm; lateral resolution, 140 μm) to visualize cryogels. The cryogel injection site was shaved and hair was removed to avoid unnecessary scattering. The scanner was placed directly above the injection site. Ultrasound images were acquired every 0.2 mm in the axial plane throughout the cryogels. After imaging, animals were allowed to recover to their normal behavior. 3D volume reconstruction of the cryogels including ectopic mineralization was performed using the multi-slice method in Vevolab software (Vevolab 5.7.1). The volume of cryogels and ectopic mineralization was created by segmenting a series of contours from 2D slices of the ultrasound images and rendering them into a 3D image. The ectopic mineralization score was performed by manual observation based on the mineralization area within the cryogels.

### Micro-Computed Tomography (microCT) acquisition and analysis

To quantify the calcified volume, we conducted microCT analysis through the Preclinical Imaging and Testing Facility at MIT. Mineralized cryogels were extracted from the flank of mice subcutaneously injected with BMP-2-loaded or blank cryogels and submerged in 4% paraformaldehyde (PFA) for fixation. After fixation, the cryogels were transferred into 70% ethanol for microCT analysis. ***MicroCT acquisition***. MicroCT imaging was performed using a Skyscan 1276 (Bruker Biospin Corp, Billerica, MA) with Skyscan acquisition software (version 1.8) and reconstructed using NRecon (version 1.7.4.2). Ex vivo samples were imaged in air within a sealed container containing ethanol-saturated sponges to minimize evaporation and prevent shrinkage of the soft phase (cellular content and unoccupied polymer), which would otherwise desiccate during scanning. Acquisition parameters were as follows: x-ray source at 55 kVp and 200 µA, 0.25 mm aluminum filter, 680 projections with a 0.3° angular step size, 315 ms exposure time, and a 2×2 detector binning mode, yielding an isotropic voxel size of 10.0 µm. ***MicroCT analysis.*** Bone and soft phase volumes were quantified from grayscale 3D microCT reconstructions. Intensity thresholds were manually selected based on prior knowledge of the material composition to differentiate bone from the soft phase. The upper threshold defined bone, while a lower threshold identified the soft phase. Both threshold values were applied to all scans. Automated methods, such as Otsu’s thresholding, did not provide biologically meaningful segmentation. Volumes were determined by summing the number of voxels assigned to each phase. Image processing and analysis were conducted using ImageJ 1.54f. The volume of cryogels (mm^3^) and the volume of calcified regions (mm^3^), thickness of trabecular (Tb.Th, cm), and bone mineral density (BMD, g/HACC) were measured from the microCT analysis. The percentage of mineralization was calculated as follows: (V_calcified_ / V_total volume_)*100. The 3D rendering of cross-sectioned tomography imaging was created by segmenting a series of contours from 2D slices of microCT images.

### Histology and immunofluorescence analysis

For the histological analysis, extracted cryogels were fixed in 4% PFA and transferred into 70% ethanol for staining. PFA-fixed cryogels were partially decalcified and embedded in paraffin wax. Sections (5 um) of the samples were stained with hematoxylin and eosin (H&E) or Masson’s Trichrome. The sections were imaged with a Nikon E800 or ZEISS Axio Scan.Z1 Slide Scanner in bright-field mode.

Immunohistochemistry was performed in paraffin sections. Paraffin sections were deparaffinized and then processed with antigen retrieval treatments using Dako Target Retrieval Solution (Agilent), prior to standard immunostaining procedures. The sectioned samples were stained using standard immunohistochemistry protocols. The samples were permeabilized with phosphate buffered saline (PBS) containing 0.1% Tween 20 or Triton X (PBST), and blocked with PBST containing 10% bovine serum albumin and 10% goat serum. The following antibodies and reagents were used for immunohistochemistry: CD45 (ab10558, Abcam), NICD (ab52301, Abcam), DLL4 (AF1389, R&D systems), Hoechst 33258 (H21491, Invitrogen), and Prolong Gold antifade reagent (Invitrogen). The sections were imaged with a ZEISS Axio Scan.Z1 Slide Scanner in bright-field mode. ImageJ was used for post-image analysis

### Flow cytometry analysis

Cells harvest was performed on long bones (femurs) and mature cryogels using Digestion buffer comprised of RPMI1640, 10% FBS, 1% Pen/Strep, 0.2mg/ml Alginate Lyase (Sigma #A1603) and Collagenase Type IV 250U/ml (Gibco # 17104019). Bone/Cryogels were chopped into pieces <1mm using a pair of scissors in presence of 2ml Digestion buffer in a 10mm petri dish. Contents are transferred into a 15ml conical tube. Petri dish is washed twice with 2ml Digestion buffer to transfer any remaining material/cells. Contents in the 15ml conical tube (6ml) are incubated in a shaker at 37°C, 200 RPM for 1 hour. The contents of 15ml tube are transferred to a 50ml tube through a 70uM cell strainer. 10ml of 1X PBS are added to the 15ml tube to rinse any stuck cells, transferred back into 50ml tube through 70uM cell strainer. Contents are spun down (500g, 4°C, 5mins). Supernatant is discarded. Samples are incubated in 500ul of ACK lysis on ice to get rid of RBCs. 10ml of FACS buffer (1x PBS + 2%FBS + 1% Pen/Strep) is added for wash, spun down (500g, 4°C, 5mins) and supernatant is discarded. Cells are transferred into a 1ml Eppendorf tube, FACS buffer added to make volume up to 1ml. Samples are now ready for Flow cytometric analyses.

Anti-mouse antibodies to CD45 (104), CD4 (GK1.5), CD8α (53-6.7), CD3ε (17A2), B220 (RA3-6B2), CD11b (M1/70), Lineage cocktail (17A2; RB6-8C5; RA3-6B2; Ter-119; M1/70), CD117 (2B8), Sca-1 (D7) CD127 (A7R34), CD150 (TC15-12F12.2), CD48 (HM48-1) and Live/Dead blue stain kit (Invitrogen # L34962), and the corresponding isotype control antibodies were purchased from BioLegend. All cells were gated based on forward and side scatter characteristics to limit debris, including dead cells. Antibodies were diluted according to the manufacturer’s suggestions. Cells were gated based on fluorescence-minus-one controls, and the frequencies of cells staining positive for each marker were recorded. To quantify T, B and myeloid cells, blood samples underwent lysis of red blood cells and were stained with anti-CD45, anti-B220, anti-CD3, anti-CD4, anti-CD8 and anti-CD11b antibodies; absolute numbers of T, B and myeloid cells were calculated using flow cytometry frequencies. Flow Cytometry data was analyzed using FlowJo.

### Statistical Analysis

All statistical analyses were performed using Prism Graphpad software version 10.4.1. Statistical tests included Student’s t-test with Welch’s correction for comparisons between two groups, and two-tail one-way ANOVA with post hoc Tukey’s multiple comparison test for group comparisons. Two-way ANOVA with repeated measures was used for circulating immune cell quantification. A p-value < 0.05 was considered statistically significant. Data are presented as means, and error bars represent the standard deviation unless otherwise noted.

## Data availability

The data that support the findings of this study are available from the corresponding authors upon reasonable request.

## Acknowledgements

The authors thank Milton Cornwall-Brady for assistance with microCT analysis, Roderick Bronson, pathologist, for assistance with histology analysis, Dr. Alex Najibi for assistance with ultrasonography imaging and Dr. Ting-Yu Shih for support with polymer synthesis. Histology imaging, consultation and services were performed in the Dana-Farber/Harvard Cancer Center in the Harvard Medical School. The authors also thank the staff at the Wyss Institute for Biologically Inspired Engineering at Harvard University for providing the support needed to perform the required experiments. This work was supported by the Lightning Biotherapeutics.

## Author contributions

S.L., K.A.-B., and A.S.S. conceptualized and designed the study. S.L., K.A.-B., A.S.S., T.T., N.D., A.S., K.S., C.J., H.I., P.K., M.C., S.E., M.C.S. were conducted experiments. S.L., K.A.-B., and A.S.S. analyzed data. S.L., K.A.-B., A.S.S., D.T.S., and D.J.M. wrote and edited the manuscript. All authors have read and contributed to editing the manuscript.

