## Supplementary Figures for "Kinetics of *de novo* Bone and Bone Marrow Niche Formation with Hybrid Click Cryogels"

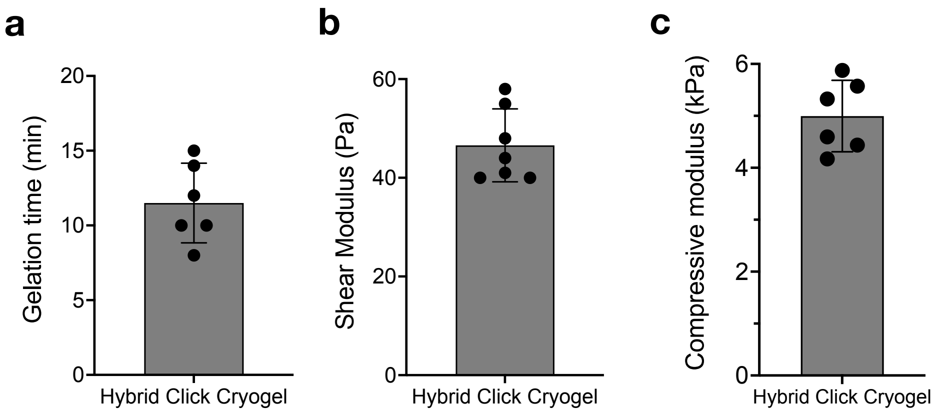


**Supplementary Figure 1. Mechanical characterization of hybrid click cryogels.** (a) Gelation time, (b) storage modulus, and (c) elastic modulus. Data represent mean ± standard deviation (SD) for n=6 cryogels per condition.

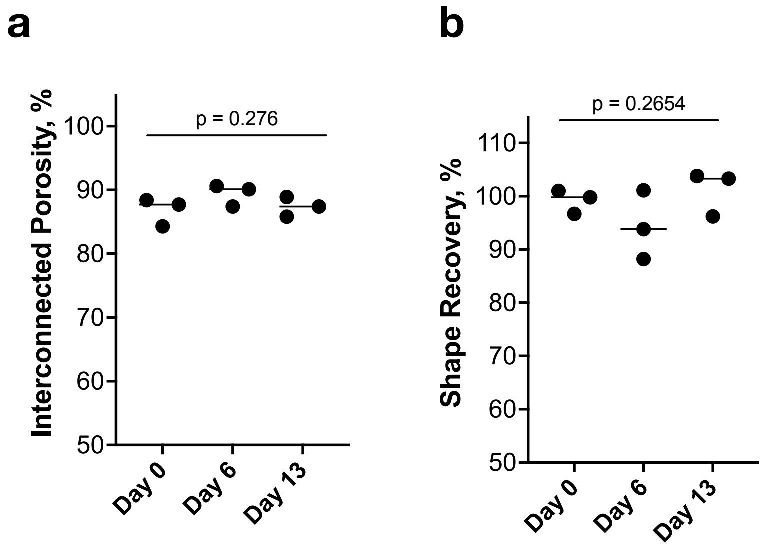


**Supplementary Figure 2.** Characterization of (a) interconnected porosity over time with incubation in tissue culture medium (DMEM with high glucose), and (b) shape recovery in hybrid click cryogels following needle injection. Data represent mean ± SD for n=3 cryogels per condition. Statistical analysis was performed via ordinary one-way ANOVA with Tukey's multiple comparison test.

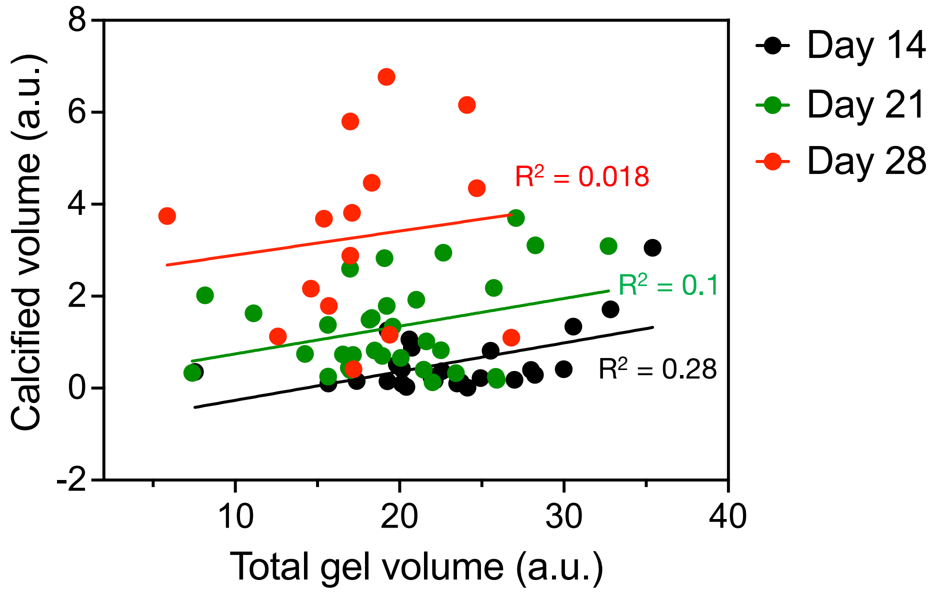


**Supplementary Figure 3.** Correlation between total cryogel volume and calcified volume over time based on ultrasonography images. Colored symbols represent individual data points, and colored lines represent correlation plots obtained through simple linear regression analysis. $R^2$ values are displayed on the plot. n= 26, 22, 15 cryogels per condition.

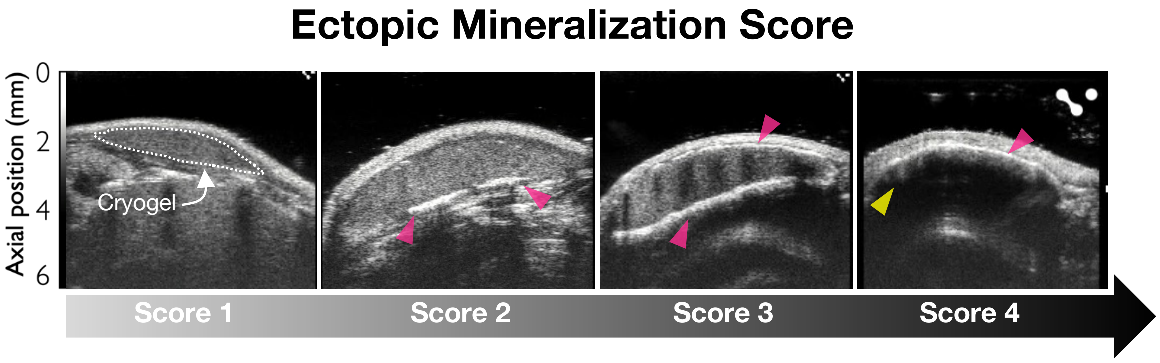


**Supplementary Figure 4.** Ultrasonography-based score metric for evaluating de novo bone formation in BMP-2 loaded cryogels implanted in subcutaneous tissue. Ectopic mineralization in cryogels appears as a dense white lining (pink arrow) with a strong shadow (yellow arrow) under ultrasonography. Representative images showing various stages of ectopic mineralization corresponding to a 4-point scoring system. 1- No mineralization, 2- Dense white scattering with a weak shadow, 3- Dense white lining with a shadow at the top or bottom and 4- Dense white lining with a strong shadow, indicating mineralization. Dashed white outline denotes the perimeter of cryogels. All ultrasonography images represent the central part of the cryogels.

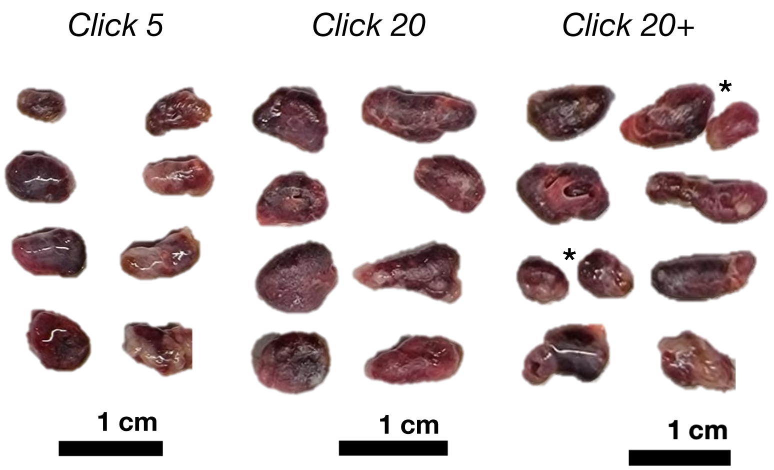


**Supplementary Figure 5.** Macroscopic images of BMP-2 loaded cryogels (day35) under different BMP-2 loading conditions. Cryogels loaded with 5 ug BMP-2 during gelation are labeled as Click 5, with 20 ug BMP-2 loaded during gelation as Click 20, and with 20 ug BMP-2 after gelation as Click 20+. Cryogels with asterisk denote those which have broken into two pieces, originating form a single cryogel.

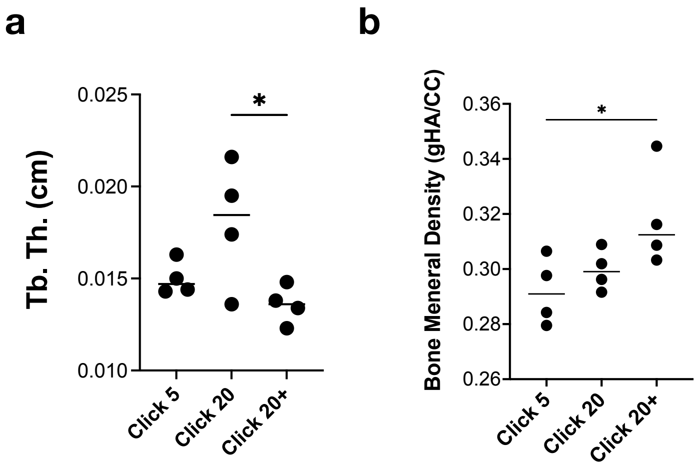


**Supplementary Figure 6. Characterization of de novo bone formation by microCT analysis.** Quantification of (a) trabecular thickness (Tb. Th.) and (b) bone mineral density in explanted cryogels (day 35). Data represent mean ± SD for n=4 cryogels per condition. Statistical analysis was performed using ordinary one-way ANOVA with Tukey's multiple comparison test. *p ≤ 0.05.

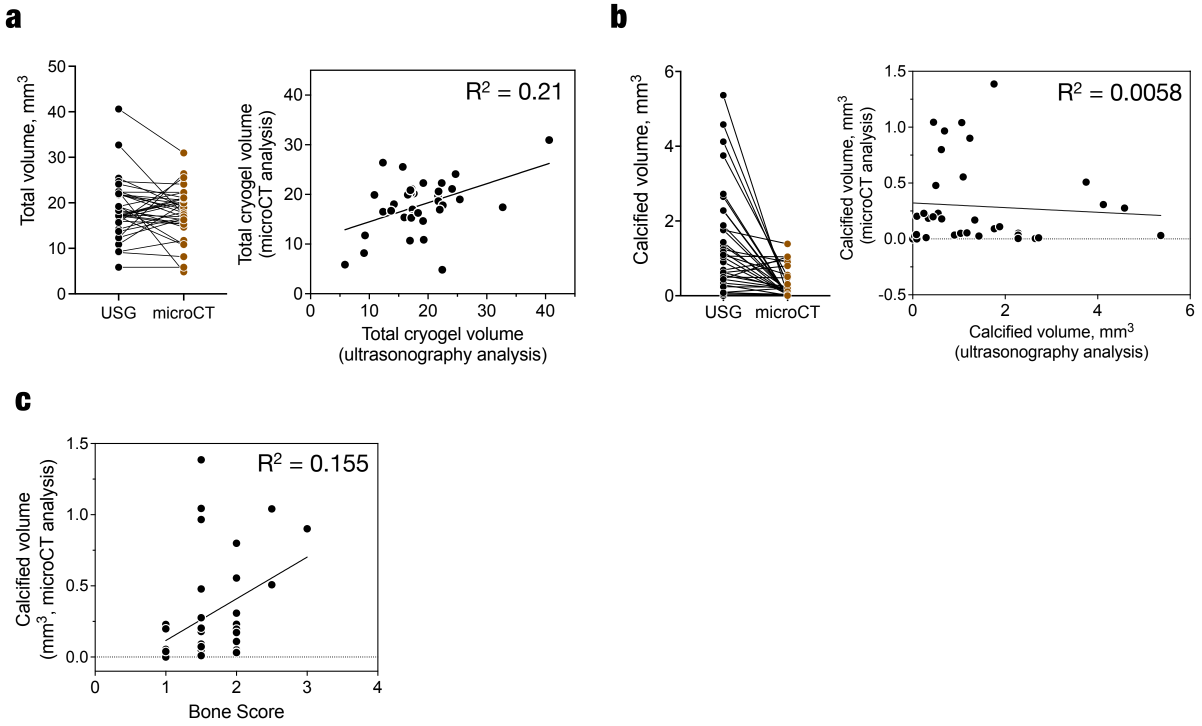


**Supplementary Figure 7.** Ultrasonographic analysis and micoCT analysis correlate in terms of total cryogel volume and Bone score. (a) Direct comparison and correlation of total cryogel volumes obtained from ultrasonography and microCT analysis. (b) Correlation of bone score obtained from ultrasonography analysis and calcified volume from microCT analysis. Simple linear regression analysis was used, and $R^2$ is denoted on each plot. n = 35 cryogels per condition.

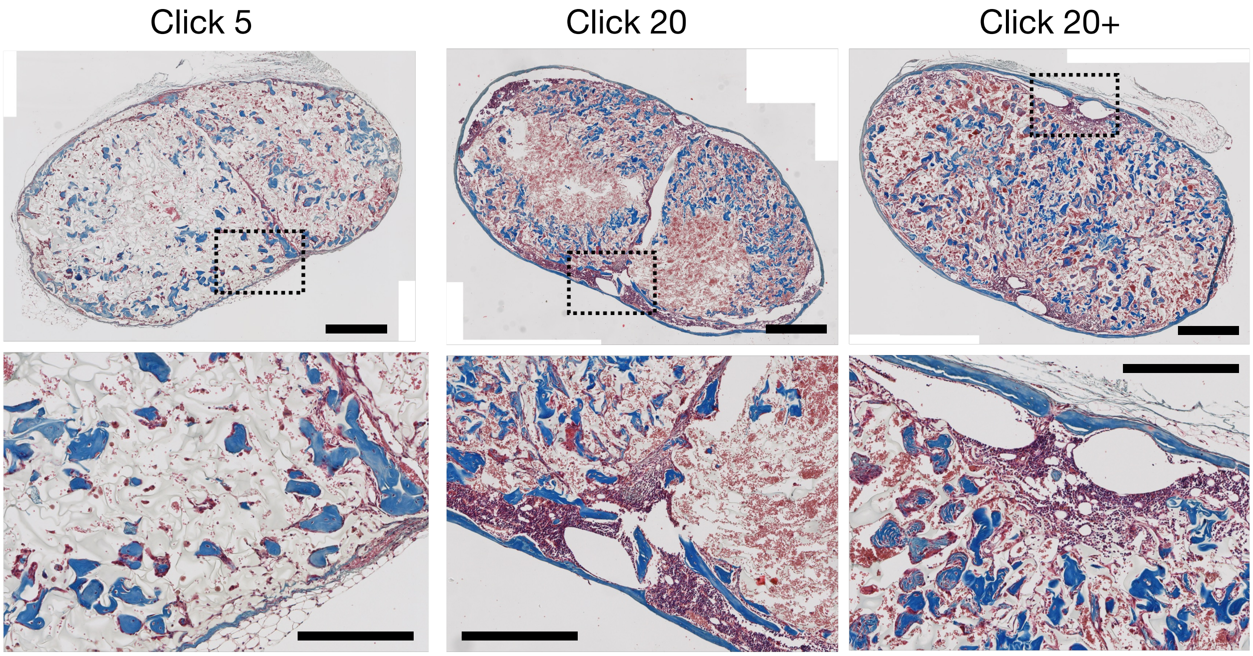


**Supplementary Figure 8.** Masson's Trichrome staining of explanted cryogels (day 35) under different BMP-2 loading conditions. Images are presented at low (top row) and high (bottom row) magnification. The bright field images show the macroporous structure of the cryogels in light-grey, with mineralization depicted in blue, located both inside and outside the cryogels. Hematopoietic tissue near the peripheral mineralization is shown in red-purple. All images represent cross-sectioned slices of paraffin-blocked cryogels. Scale bars represent 500 μm (top row), 250 μm (bottom row).

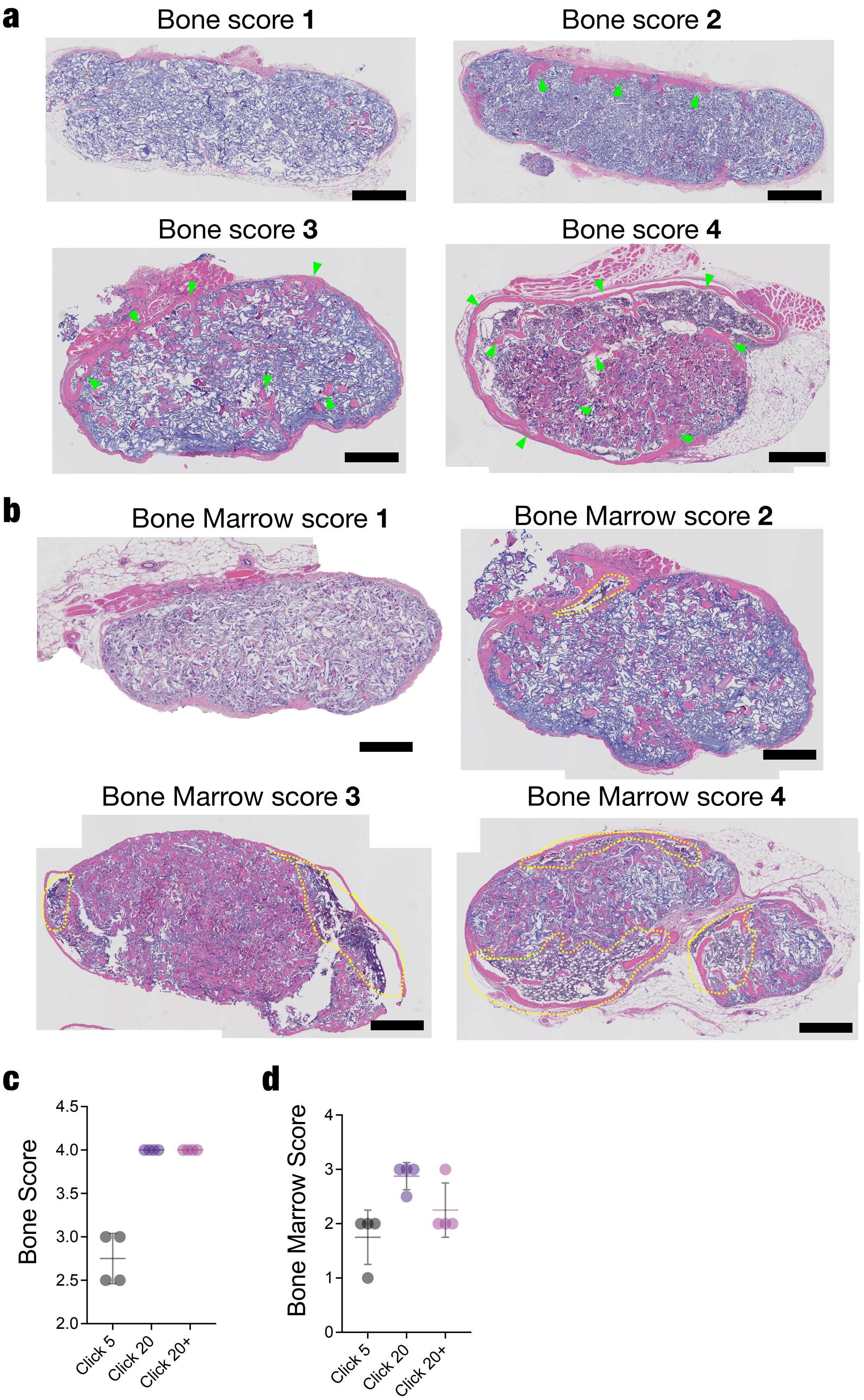
a
Bone score 1
Bone score 2
Bone score 3
Bone score 4
b
Bone Marrow score 1
Bone Marrow score 2
Bone Marrow score 3
Bone Marrow score 4
c
Bone Score
4.5
4.0
3.5
3.0
2.5
2.0
Click 5
Click 20
Click 20+
d
Bone Marrow Score
4
3
2
1
0
Click 5
Click 20
Click 20+

**Supplementary Figure 9.** Histology-based score metric for evaluating de novo bone and bone marrow-like niches formation in BMP-2 loaded cryogels (day 35). The bone and bone marrow scores are determined independently based on the relative amount of mineralization and hematopoietic tissue, respectively: 1 - no mineralization or no hematopoietic tissue; 2 - tiny spicules present or small regions of hematopoietic tissue at the peripheral region; 3 - numerous spicules with small cortical bone or medium-sized regions of hematopoietic tissue at the peripheral region; 4 – extensive cortical bone with spicules or extensive hematopoietic tissue present in either peripheral or central regions. Representative brightfield images of Hematoxylin and Eosin (H&E) stained cryogels showing different bone scores (a) and bone marrow scores (b). Cryogels are shown in light purple, with mineralization appearing in pink at the peripheral and central regions of the cryogels. Mineralization and Hematopoietic tissue is observed in peripheral or central regions labeled with green arrows and yellow dotted lines, respectively. All images represent cross-sectioned slices of a paraffin-blocked cryogels. Scale bars represent 500 µm. Comparison of (c) bone score and (d) bone marrow score under different conditions. Data represents mean ± standard deviation (n=4).

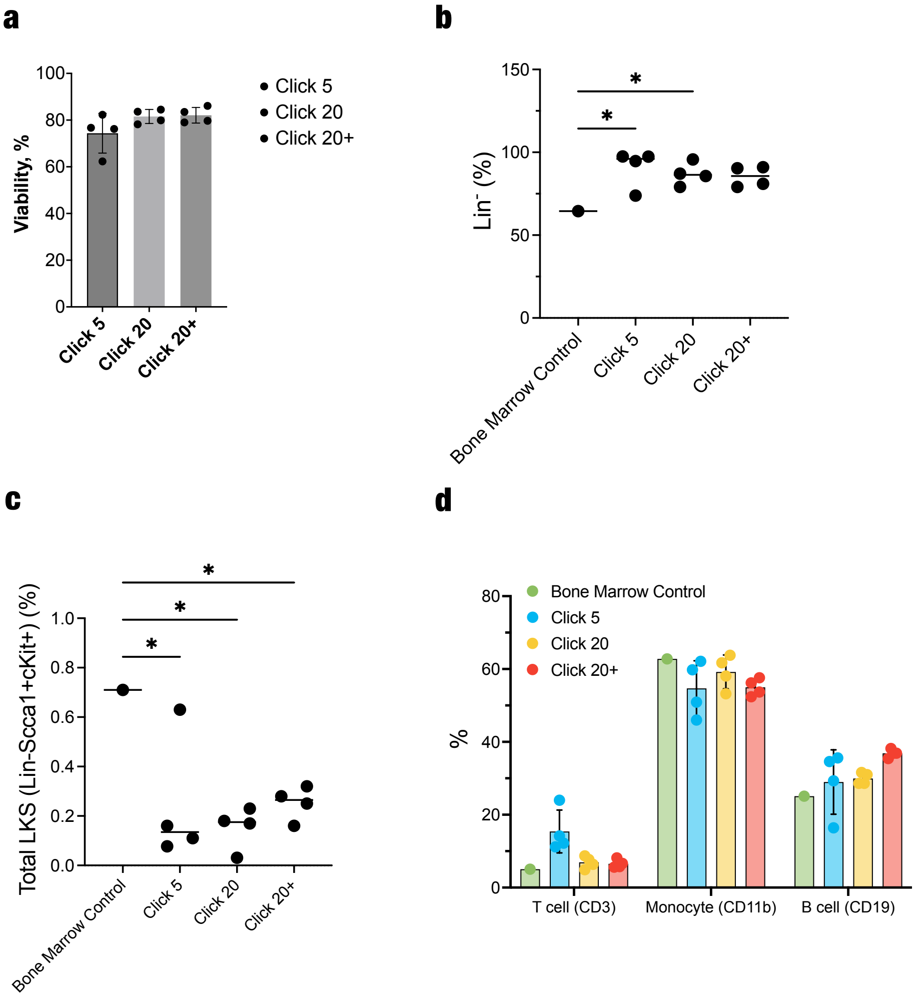


**Supplementary Figure 10.** Flow cytometric analysis of explanted cryogels under different BMP-2 loading conditions. (a) Cell viability, (b) percentage of lineage-negative cells, (c) percentage of Lin-Sca1+cKit+ (LKS) cells, (d) percentage of mature cells including T cells (CD3+), monocytes (CD11b+), B cells (CD19+) in explanted cryogels. Bone marrow from non-BMP-2 treated mice was used as a control in (b-d). Data represents mean ± standard deviation for n=4 cryogels per condition. Statistical comparison was performed using an ordinary one-way ANOVA with Fisher's LSD analysis (b, c). $^{*}p \leq 0.05$.

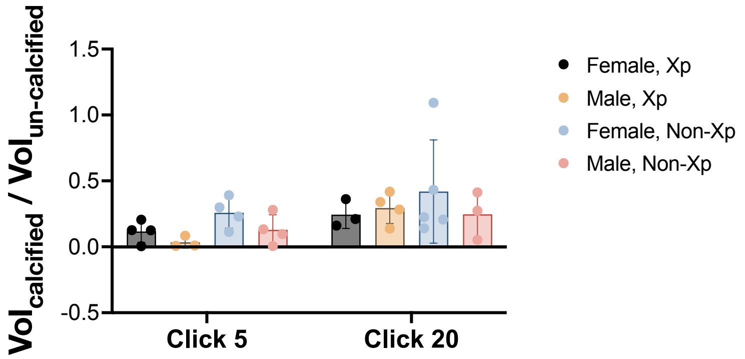


**Supplementary Figure 11.** Calcified-to-uncalcified volume ratio of explanted cryogels (day 35). Data represent mean ± SD for n=4 cryogels.

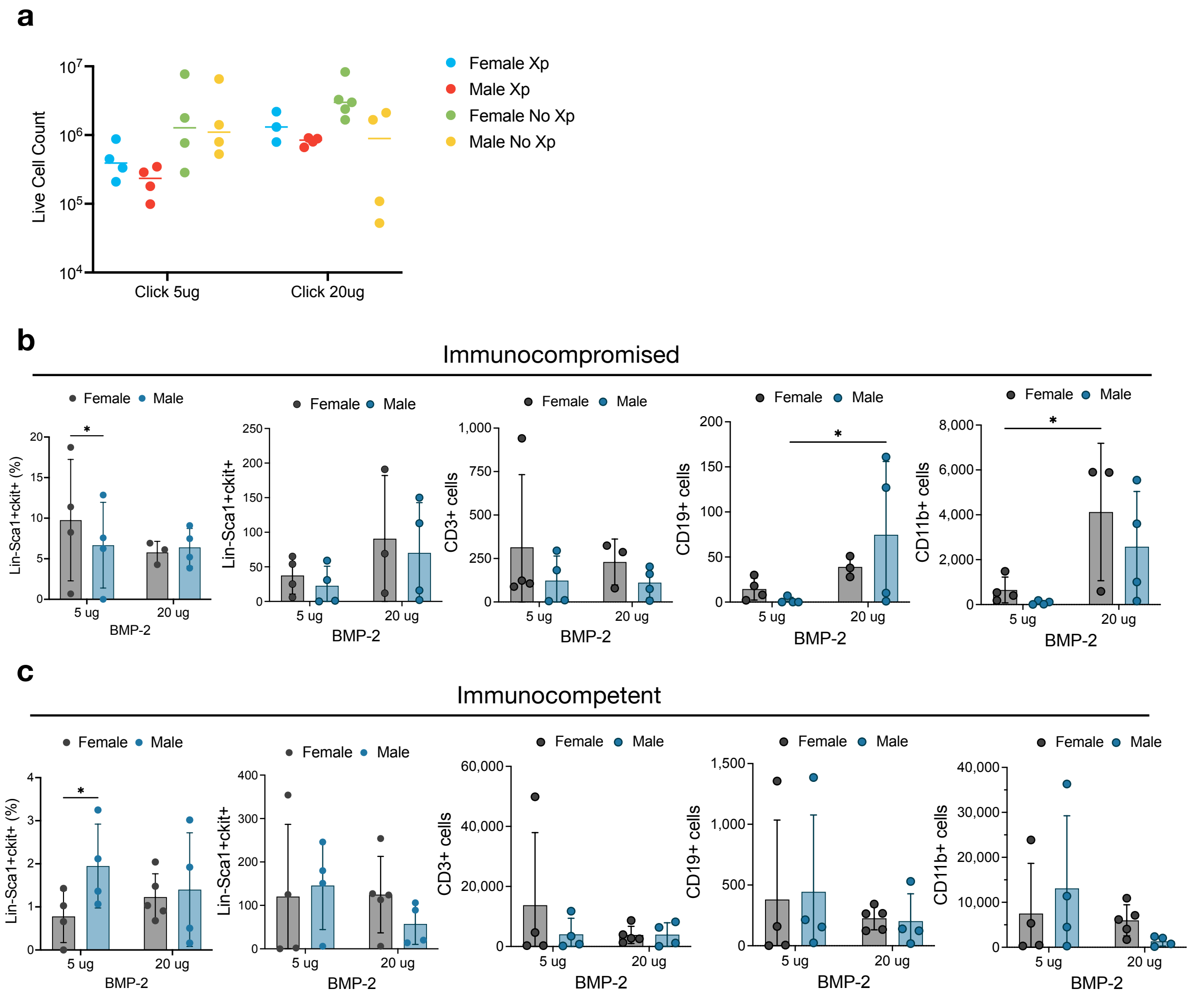


**Supplementary Figure 12.** Flow cytometric analysis of cells isolated from explanted cryogels loaded with 5 ug or 20 ug BMP-2 (day 35) under different biological sex and immunocompetency conditions. Comparisons were made between female and male mice, as well as between immunocompromised (Xp) and immunocompetent (No-Xp) conditions. (a) Number of live cells. (b, c) LKS cells, T cells (CD3+), B cells (CD19+), neutrophils (CD11b+) population in the immunocompromised (b) or in the immunocompetent conditions (c), comparing female and male mice. Data represent mean ± SD for n=4 or 5 cryogels per condition. Statistical comparison was performed using a two-way ANOVA with Fisher's LSD analysis. *p ≤ 0.05.

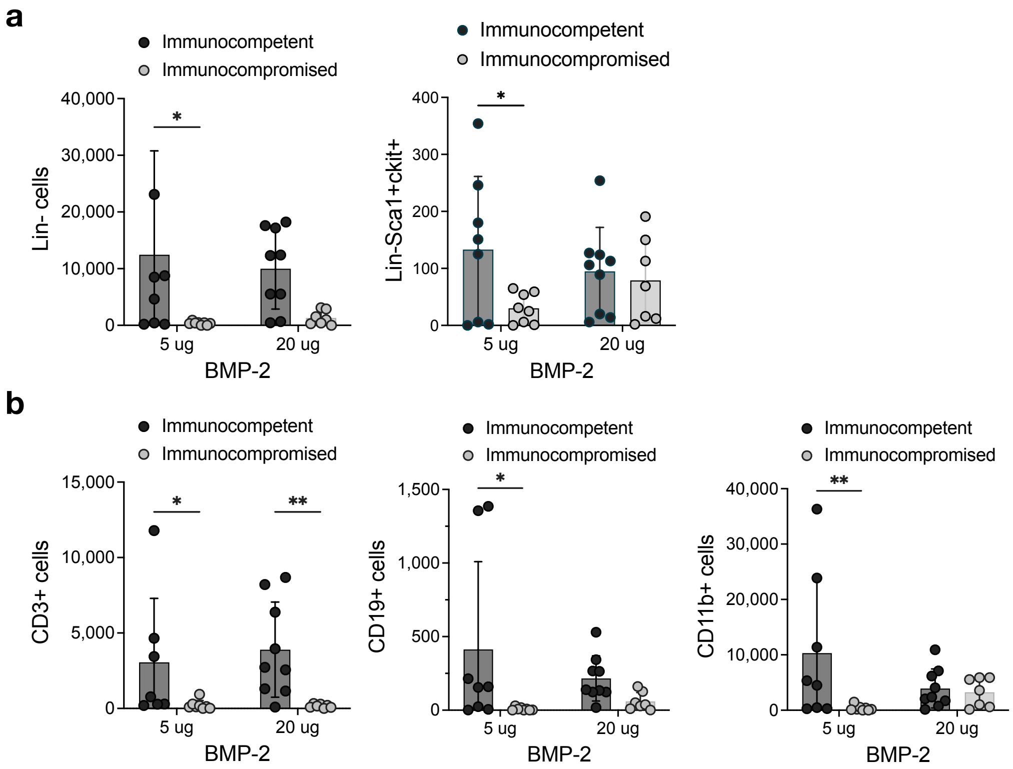


**Supplementary Figure 13.** Flow cytometric analysis of cells isolated from explanted cryogels loaded with 5 ug or 20 ug BMP-2 (day 35) under different immunocompetency conditions. (a) Comparison of lineage-negative and LKS cell populations between immunocompromised and immunocompetent conditions. Data from Figures 5G and 5H are replotted here for comparison. (b) Comparison of T cells (CD3+), B cells (CD19+), and neutrophils (CD11b+) populations between immunocompromised and immunocompetent conditions. Data represent mean ± SD for n= 8 or9 cryogels per condition. Statistical comparison was performed using a two-way ANOVA with Fisher's LSD analysis. *$p \leq 0.05$ and **$p \leq 0.01$.

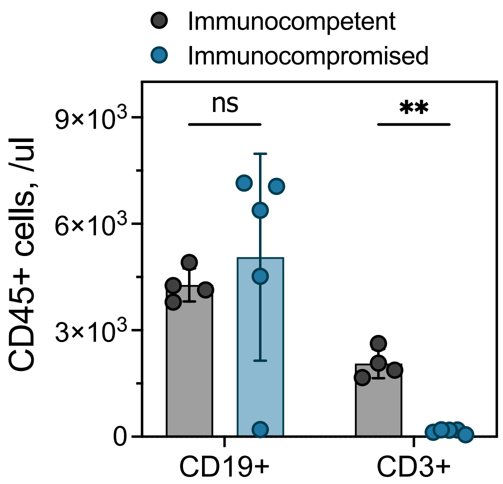


**Supplementary Figure 14.** Comparison of B cell (CD45+CD19+) and T cell (CD45+CD3+) populations in peripheral blood on day 28 between immunocompromised and immunocompetent conditions. Data represent mean ± SD for n=5 cryogels per condition. Statistical comparison was performed using multiple unpaired t-tests with Welch's correction. **p ≤ 0.01.

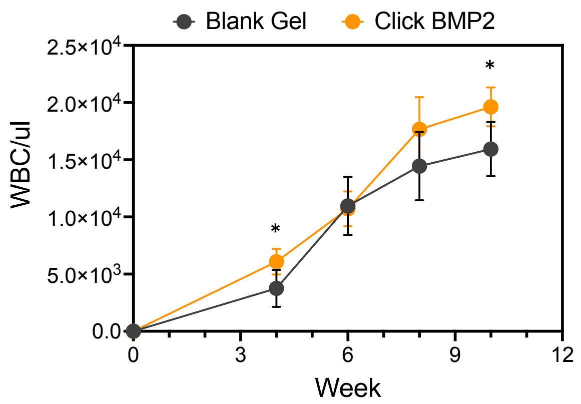


**Supplementary Figure 15.** Circulating white blood cell count over time. Data represent mean ± SD for n=7 per Blank gel and 10 cryogels per Click BMP2. Statistical comparison was performed using a two-way ANOVA with Šídák's multiple comparison test; *$p \leq 0.05$.

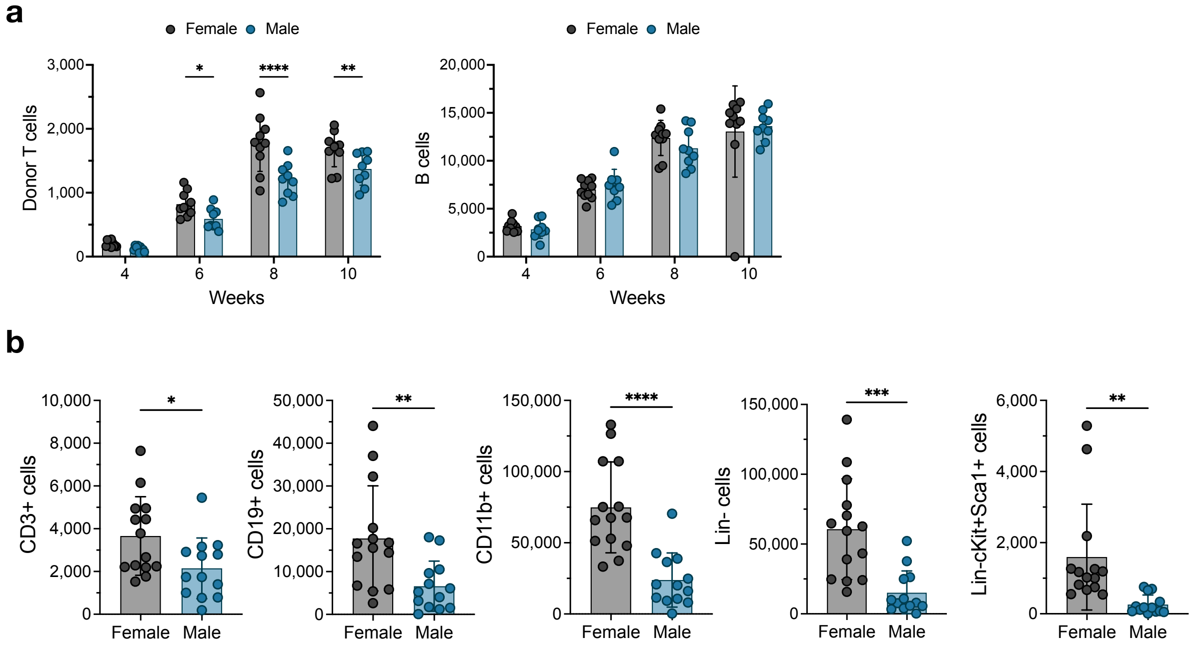


**Supplementary Figure 16.** Quantification of circulating immune cell populations and cells resided in explanted cryogels (day 84) in animals of different biological sex. (a) Comparison of circulating donor T cells and B cells between female and male mice. (b) Comparison of T cells (CD3+), B cells (CD19+), neutrophils (CD11b+), lineage-negative cells, LKS cells (Lin-cKit+Sca1+) in explanted cryogels in both female and male mice. Data represent mean ± SD for n= 10 (a), 14 (b) cryogels per condition. Statistical comparison was performed using a two-way ANOVA with Fisher's LSD analysis (a) and un-paired t test analysis (b). *$p \leq 0.05$, **$p \leq 0.01$, ***$p \leq 0.001$, ****$p \leq 0.0001$.

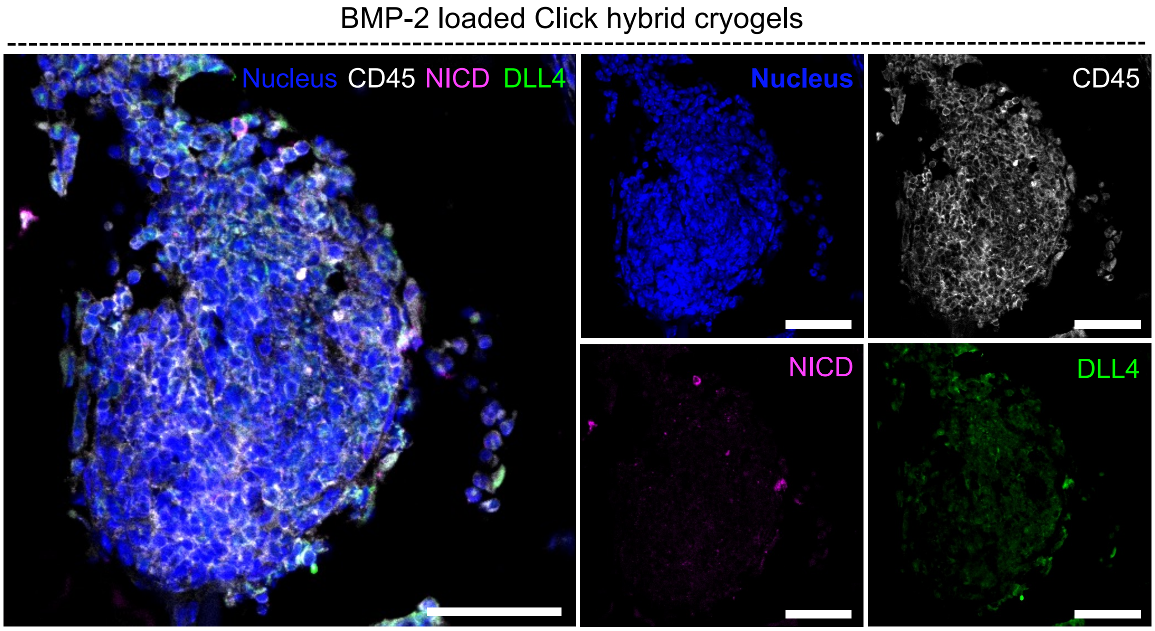


**Supplementary Figure 17.** Focused view of hematopoietic tissue region from BMP-2 loaded click hybrid click cryogel explanted at day 84 from immunocompromised mouse. Explanted cryogels were stained for CD45 (immune cells), NICD (Notch-activated cells), and DLL4. Representative fluorescent images of hematopoietic tissue regions in cryogels showing a pool of cells with evidence of DLL4/Notch signaling. Images represent cross-sectioned slices of paraffin-blocked cryogels. Scale bars represent 50 µm.
